# ALLOPURINOL BLOCKS AORTIC ANEURYSM IN A MOUSE MODEL OF MARFAN SYNDROME VIA REDUCING AORTIC OXIDATIVE STRESS

**DOI:** 10.1101/2021.10.13.464182

**Authors:** Isaac Rodríguez-Rovira, Cristina Arce, Karo De Rycke, Belén Pérez, Aitor Carretero, Marc Arbonés, Gisela Teixidò-Turà, Mari Carmen Gómez-Cabrera, Victoria Campuzano, Francesc Jiménez-Altayó, Gustavo Egea

## Abstract

**Background:** Increasing evidence indicates that redox stress participates in MFS aortopathy, though its mechanistic contribution is little known. We reported elevated reactive oxygen species (ROS) formation and NADPH oxidase NOX4 upregulation in MFS patients and mouse aortae. Here we address the contribution of xanthine oxidoreductase (XOR), which catabolizes purines into uric acid and ROS in MFS aortopathy.

**Methods and Results:** In aortic samples from MFS patients, XOR protein expression, revealed by immunohistochemistry, increased in both the tunicae intima and media of the dilated zone. In MFS mice (*Fbn1^C1041G/+^*), aortic *XOR* mRNA transcripts and enzymatic activity of the oxidase form (XO) were augmented in the aorta of 3-month-old mice but not in older animals. The administration of the XOR inhibitor allopurinol (ALO) halted the progression of aortic root aneurysm in MFS mice. ALO administrated before the onset of the aneurysm prevented its subsequent development. ALO also inhibited MFS-associated endothelial dysfunction as well as elastic fiber fragmentation, fibrotic collagen remodeling, nuclear translocation of pNRF2 and increased 3’-nitrotyrosine levels all occurring in the tunica media. ALO reduced the MFS-associated large aortic production of H_2_O_2_, and NOX4 and MMP2 transcriptional overexpression.

**Conclusions:** Allopurinol interferes in aortic aneurysm progression acting as a potent antioxidant. This study strengthens the concept that redox stress is an important determinant of aortic aneurysm formation and progression in MFS and warrants the evaluation of ALO therapy in MFS patients.

**HIGHLIGHTS:**

- Xanthine oxidoreductase (XOR) is upregulated in the aortic aneurysm of Marfan syndrome (MFS) both in patients and young mice.
- Allopurinol halts the formation and progression of aortic aneurysm in MFS mice
- Allopurinol reduces a variety of oxidative stress-associated molecular reactions.
- Allopurinol prevents MFS endothelial-dependent vasodilator dysfunction.
- The antioxidant action of allopurinol suggests its repositioning for pharmacological use in MFS aortopathy.

**GRAPHICAL ABSTRACT:** 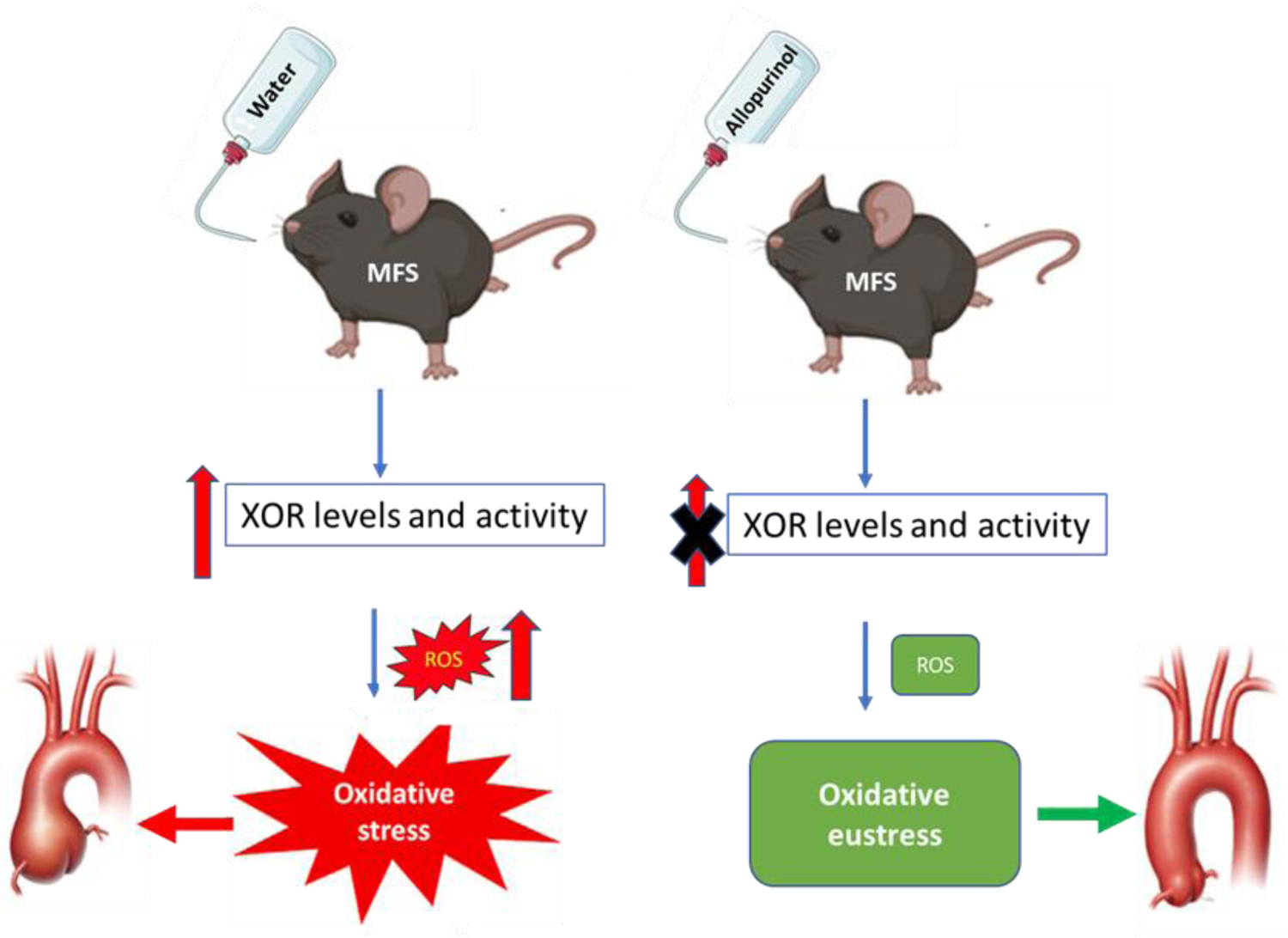

## INTRODUCTION

Marfan syndrome (MFS) is a relatively common inherited rare disease of the connective tissue caused by mutations in the gene encoding the extracellular matrix glycoprotein fibrillin-1^1^. This multisystem disease mainly affects the skeleton (abnormally long bones and spine deformations), eyes (lens dislocation) and aorta (aortic root aneurysm). The latter commonly leads to aortic dissection and rupture, the main cause of reduced life expectancy in patients^2^.

Reactive oxygen species (ROS) consist of radical (anion superoxide/O_2_^-^ and hydroxyl radical /HO·) and non-radical (hydrogen peroxide/H_2_O_2_) oxygen species, which are highly reactive chemicals formed by the partial reduction of oxygen. O_2_^-^ and H_2_O_2_ are the most frequently formed ROS and, under physiological concentrations, have important signaling functions (oxidative eustress)^3^. However, when ROS overwhelm the intrinsic cellular antioxidant system, either via an abnormal overproduction of ROS or reduction of their antioxidant capacity, they contribute to pathogenesis (oxidative stress), causing transient or permanent damage to nucleic acids, proteins, and lipids^4, 5^.

In the cardiovascular system, ROS are involved in, among other dysfunctions, the development of aortic abdominal aneurysms (AAA) through the regulation of inflammation induction, matrix metalloproteinases (MMPs) expression, vascular smooth muscle cell (VSMC) apoptosis and phenotypic changes and modifying extracellular matrix properties^6–10^. However, the impact of ROS on thoracic aortic aneurysm (TAA) of genetic origin, such as in MFS and Loeys-Dietz syndrome (LDS), is less known^11^. We and others have previously shown that ROS production is increased in MFS^12–18^, although the specific generators of ROS and their respective impact on the formation and/or progression in either human or murine MFS aortic aneurysms is still poorly understood^19, 20^. We reported the involvement of upregulated NADPH (nicotinamide adenine dinucleotide phosphate) oxidase 4 (NOX4) both in human and mouse MFS aortic samples and cultured VSMCs^17^. Nonetheless, besides NOX4, another important source of ROS in the cardiovascular system is xanthine oxidoreductase (XOR)^21^.

XOR is involved in the final steps of nucleic acid-associated purine degradation and, particularly, in the conversion of hypoxanthine into xanthine and xanthine into urate. XOR exists in two forms that are derived from a single gene (*XDH*)^22^. The reduced form of XOR is referred to as xanthine dehydrogenase (XDH), and the oxidized form as xanthine oxidase (XO). XDH can be post-translationally modified to XO via proteolysis or oxidation of critical cysteines^23^. The XDH form has greater abundance and affinity for NAD^+^ as the electron acceptor to generate NADH, which is a critical substrate for NADPH oxidases. Likewise, the XO form is mainly associated with the production of large amounts of O_2_^-^ and H_2_O_2_ by preferentially using oxygen as the electron acceptor^24, 25^. Under healthy conditions, XDH is constitutively expressed, and XO levels are low both in plasma and heart^26^. However, XDH conversion into XO is favored by hypoxia, low pH, ischemia, inflammation, and the presence of peroxynitrite and H_2_O_2_ itself^27–29^. XOR is widely distributed throughout the organism mainly in the liver and gut, but also present in intestine, lung, kidney, myocardium, brain and plasma^30^. In the vascular endothelium, XOR is bound to cell surface glycosaminoglycans^26^; the enhancement of endothelium attached XOR favors local ROS production with subsequent endothelial dysfunction^31^. XOR-derived anion superoxide (O_2_^-^) easily interacts with endothelial cell-generated nitric oxide (NO) forming peroxynitrite (ONOO^-^), which in the endothelium of the tunica intima and VSMCs of the media irreversibly generates reactive nitrogen species (RNS) residues such as 3’-nitrotyrosine^32, 33^, which is usually used as a redox stress marker. Increased XOR levels have been reported in aortic samples from adult MFS mice^14^ and LDS patients^34^, even though the extent to which XOR contributes to TAA pathogenesis is unknown.

On the other hand, concomitantly to ROS production, XOR generates uric acid (UA), which in humans can pathologically accumulate in the plasma and some tissues. In rodents, UA is rapidly catabolized by uricase (absent in humans) to allantoin^35^. UA has a dual role in redox biology, acting as an antioxidant (both *in vitro* and *in vivo*) accounting for as much as 50% of the total antioxidant capacity of biological fluids in humans^36^. However, when UA accumulates in the cytoplasm or in acidic/hydrophobic milieu, it acts as a pro-oxidant promoting redox stress^36, 37^. UA was found in the wall of human aortic aneurysms and atherosclerotic arteries^38^ and there was a positive correlation between serum UA levels and aortic dilation and dissection^39, 40^. Epidemiologic and biochemical studies on UA formation have demonstrated that it is not only UA itself that leads to a worse prognosis and increased cardiovascular events, but also ROS formed during XOR activity. Therefore, the resulting combined action of excessively formed UA and ROS surely contribute to oxidative stress-linked cardiovascular pathological events^41^.

The aim of this study was to investigate the participation of XOR in MFS aortopathy in more detail and evaluate the effectiveness of the XOR inhibitor allopurinol (ALO) on the formation and/or progression of MFS-associated aortopathy. ALO is an analogue of hypoxanthine and, therefore, a competitive inhibitor of XOR^42^; it is a drug routinely prescribed to treat hypouricemic and hypertensive patients^43^. XOR also functions beyond its basic housekeeping role in purine catabolism, acting in oxidant signaling, which could contribute to exacerbating oxidative stress-associated MFS aortopathy. Therefore, since XOR is an important source of ROS in the cardiovascular system, we hypothesized that, together with upregulated NOX4^17, 44^, the formation and/or progression of aortic aneurysm in MFS is favored by abnormal aortic XOR activity with the consequent dysfunctional increase in ROS levels (and likely UA and/or allantoin as well) exacerbating the oxidative stress-associated MFS aortopathy.

## METHODS

Please see Methods in Supplemental Materials.

Please see the Major Resources Table in Supplemental Materials.

## RESULTS

### XOR is augmented in the dilated aorta of MFS patients

We evaluated protein levels by immunohistochemistry with specific anti-XOR antibodies in the aorta of MFS patients subjected to aortic reparatory surgery. Both the tunicae intima and media of the dilated aortic zone presented a significant increase in immunostaining compared with adjacent non-dilated and healthy aortae (Fig. 1), which is demonstrative of an increased protein expression of XOR associated with aortic aneurysm in MFS patients. No correlation between the intensity of the immunostaining and the age of the patients analyzed was observed.

**Figure 1.**
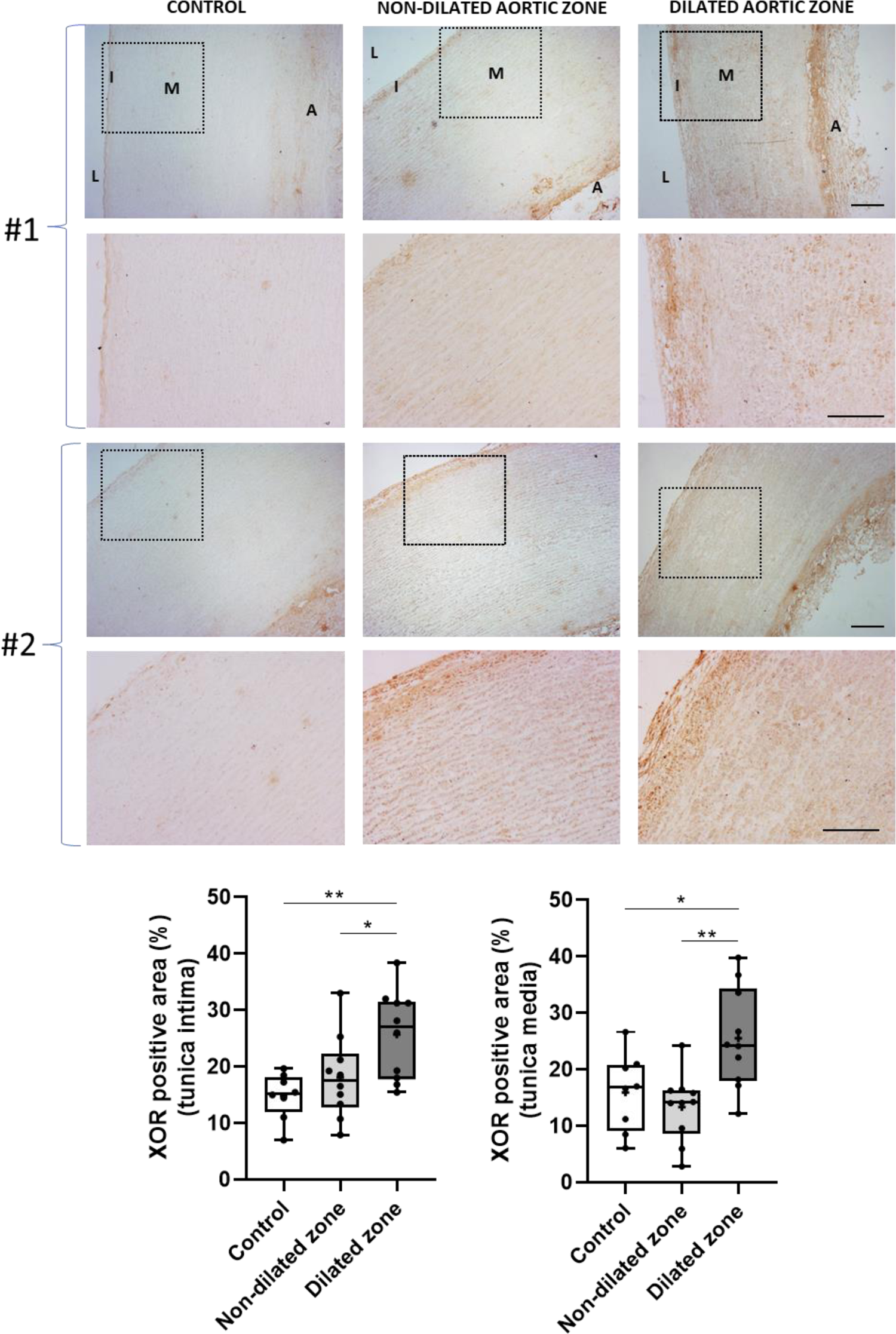
Xanthine oxidoreductase is upregulated in the aorta of MFS patients. Representative images of XOR immunostaining in the ascending aorta from two healthy heart donors (control) and in the dilated and adjacent non-dilated aneurysmal zones of the aorta from two representative MFS patients (#1 and #2). The lower panels of each aortic sample are magnified images from the corresponding upper sample (black squares). L=lumen; I: tunica intima; M: tunica media; A: tunica adventitia. Quantitative analysis of immunolabeling results evaluated in tunicae intima and media. Bars, 100 µm. Results are the mean±SEM (n=8-10). Statistical test analysis: One-way ANOVA; *p≤0.05 and **p≤0.01.

### XOR mRNA expression and enzymatic activity are only increased in the aorta of young MFS mice

We next evaluated whether XOR was also increased in the MFS mouse aorta. Firstly, we evaluated XOR transcriptional levels by RT-PCR in 3- and 6- and 9-month-old WT and MFS mice. Aortae from 3-month-old MFS mice showed significantly increased levels of *XOR* transcripts compared with age-matched WT animals. This increase was not observed in 6- or 9-month-old mice (Fig. 2A). Immunohistochemistry with anti-XOR antibodies confirmed the increased expression of XOR in both the intima and media tunicae only in 3-month-old MFS mice aortae (Fig. 2B).

**Figure 2.**
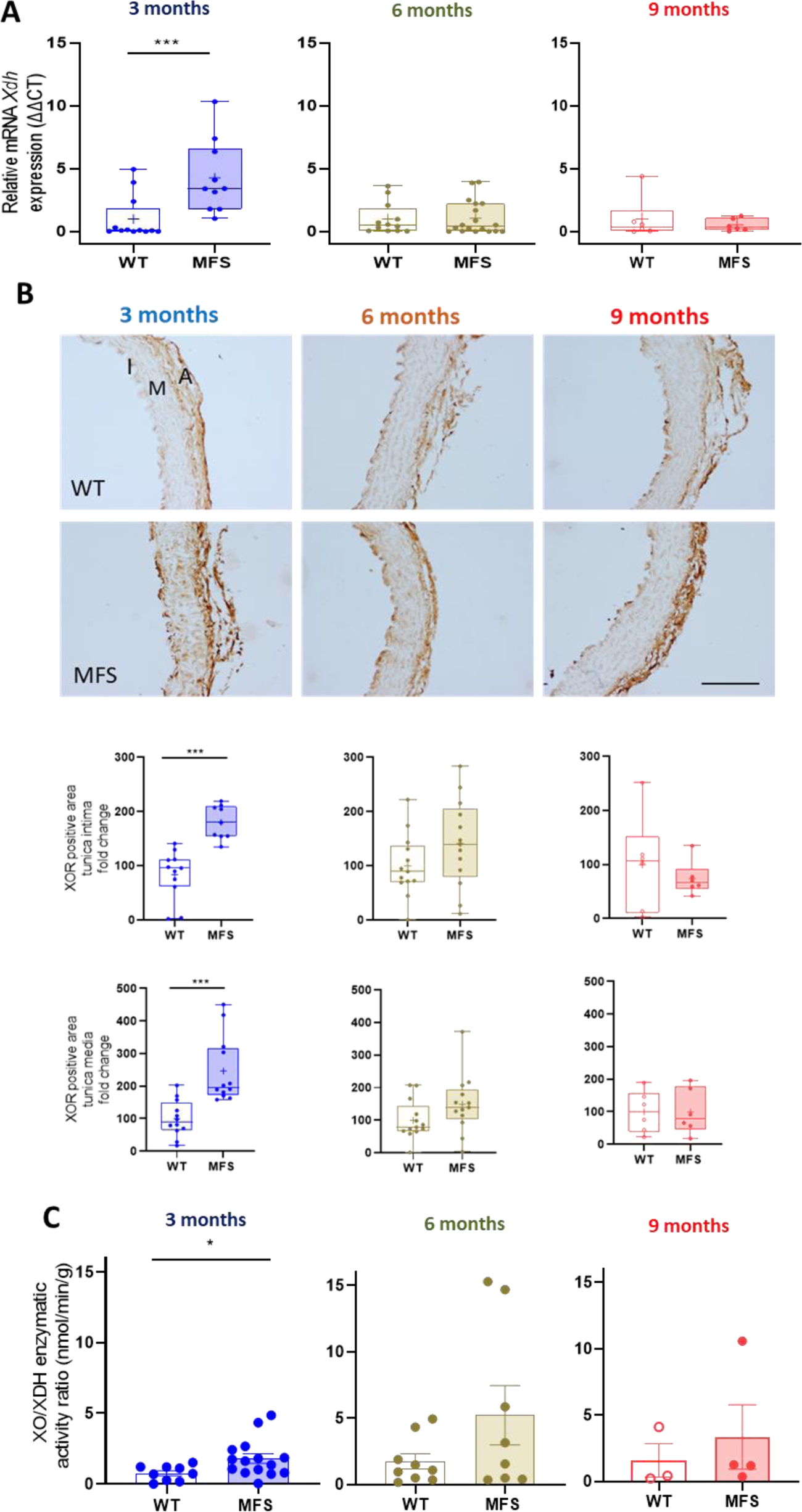
XOR expression levels and enzymatic activity in the MFS mouse ascending aorta. **(A)** mRNA expression levels of XOR (*Xdh*) in WT and MFS mice of different ages (3-, 6- and 9-month-old; n=10-15). **(B)** XOR protein levels revealed by immunohistochemistry with anti-XOR antibodies in paraffin-embedded aortae from 3-, 6- and 9-month-old WT and MFS mice. Below of images, the quantitative analysis of the respective HRP immunostaining in tunicae intima and media. I: tunica intima; M: tunica media; A: tunica adventitia. Bar, 100 µm. **(C)** XO/XDH enzymatic activity ratio in WT and 3- and 6-month-old MFS mice. Data represented as boxplots. Statistical test analysis: Mann Whitney U test (A-C). ***p ≤0.001 and *p ≤0.05.

To differentiate the functional contribution of the XDH and XO protein forms of XOR in the MFS aorta, we next measured their respective enzymatic activities in aortic lysates (see a representative assay in Fig. S2). The XO/XDH enzymatic activity ratio was significantly higher in 3-month-old but not in 6- or 9-month-old MFS mice (Fig. 2C), which is in accordance with transcriptional results (Fig. 2A). In this manner, results complement the previously reported upregulation of XOR in MFS mouse aortae.^14^

### Allopurinol inhibits the progression of aortic aneurysm and prevents its formation in MFS mice

The upregulation and activity of XOR in MFS aortae suggest its contribution to oxidative stress in MFS aortopathy. We hypothesized that the inhibition of its activity could ameliorate aortic aneurysm progression. To test this hypothesis, we treated WT and MFS mice with ALO. We first palliatively treated 2-month-old mice with ALO until 6 months of age (PA1; Fig. S1) and then evaluated aneurysm progression by ultrasonography. ALO significantly reduced the characteristic enlarged aortic root diameter occurring in MFS mice, the diameter obtained being highly similar to WT animals (Fig. 3A and Table S2). ALO did not cause any alteration in the aortic root diameter of WT mice. Moreover, no sex differences were observed regarding the effectiveness of ALO (Table S2).

**Figure 3.**
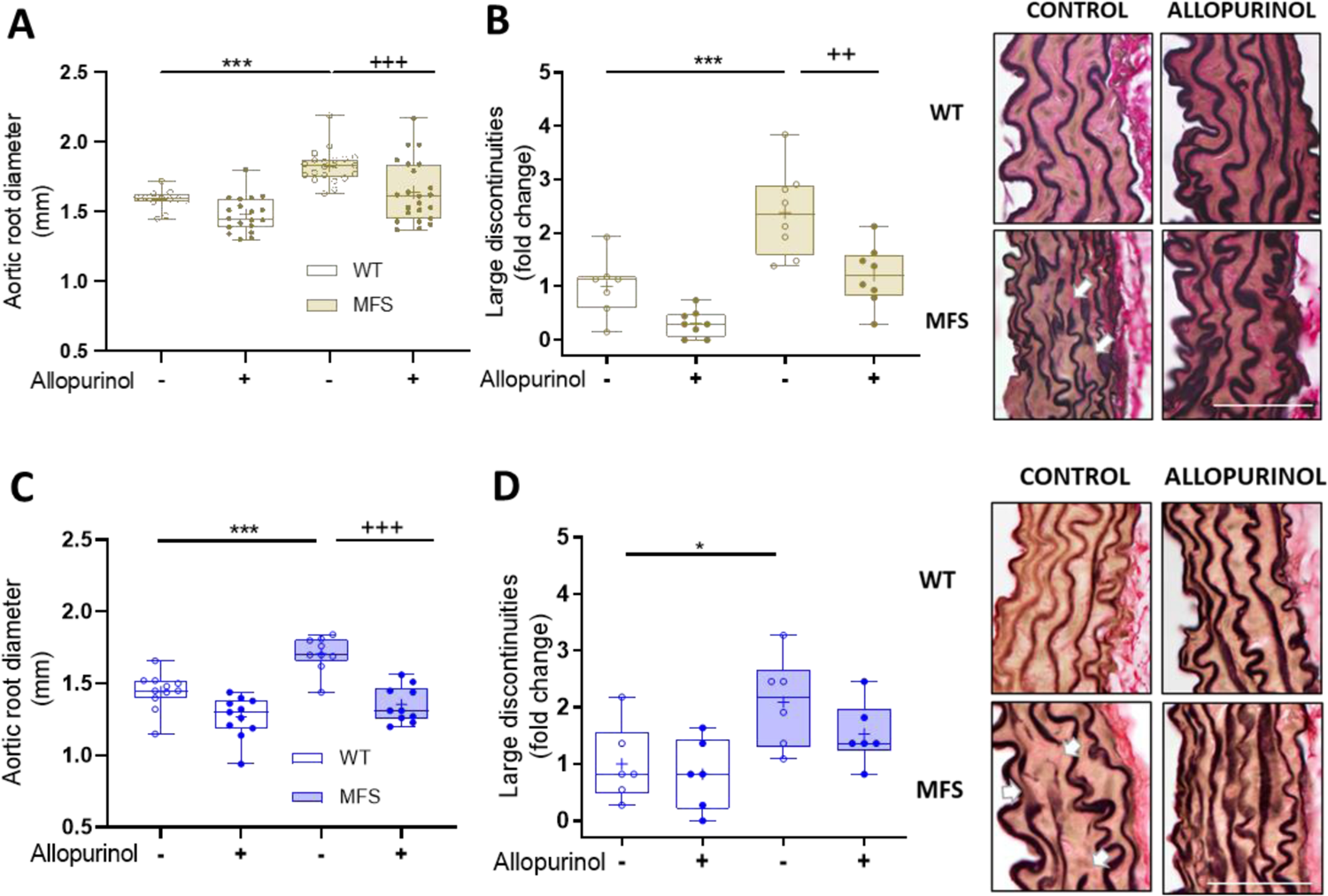
Allopurinol prevents both the formation and progression of aortic root dilation in MFS mice. **(A)** Allopurinol halts the progression of aortic root dilatation in MFS mice measured by ultrasonography in WT and MFS mice of 6 months of age treated with allopurinol for 16 weeks (PA1). See also Table S2. (**B)** The number of large discontinuities in the elastic lamina of the aortic media (360°) from WT and MFS mice palliatively treated with allopurinol (n=14-16). **(C)** Allopurinol prevents the formation of aortic root dilatation in MFS mice measured by ultrasonography in 3-month-old WT and MFS mice preventively treated with allopurinol for 12 weeks (PE). See also Table S3. **(D)** Number of large discontinuities in the elastic lamina of the aortic media (360°) from WT and MFS mice preventively treated with allopurinol (n=9-11). On the right of panels B and D, representative Elastin Verhoeff-Van Gieson staining in paraffin sections of the ascending aorta. White arrows in B and D indicate examples of the large discontinuities analyzed. Bar, 100 µm. Data represented as boxplots. Statistical test analysis: Two-way ANOVA and Tukey post-test. ***^/+++^*P*≤0.001, ^++^*P*≤0.01 and \**P*≤0.05; *effect of genotype; ^+^effect of treatment.

We also evaluated aortic wall organization by histomorphology, quantifying the number of large elastic fiber ruptures as previously defined^45^. ALO treatment in MFS mice caused an apparent normalization of elastic fiber organization and integrity, their overall morphology being indistinguishable from non-treated WT animals (Fig. 3B).

ALO has also been safely administered during the late stages of pregnancy, demonstrating efficiency in reducing uric acid, hypertension, and oxidative stress levels^46–49^. Therefore, we next evaluated whether ALO also prevented the formation of the aneurysm. ALO was administered to pregnant and lactating female mice and then to weaning offspring until the age of 3 months (PE; Fig. S1). ALO fully prevented aneurysm formation (Fig 3C and Table S3), also showing a trend to reduce elastic fiber breaks (Fig. 3D).

We also analyzed the diameter of the tubular portion of the ascending aorta. In the PA1 treatment, ALO also normalized the diameter of MFS aortae (Fig. S3A; Table S4). However, this was not the case for the PE treatment (Fig. S3B; Table S5), likely because of the shorter ALO treatment and the younger age of the MFS mice treated, in which the ascending aorta was still not sufficiently affected as in the PA1 treatment.

### The inhibition of aortopathy by allopurinol is reversible after drug withdrawal

We next examined whether the progression of the aortopathy inhibited by ALO was permanent or transient. To this end, we evaluated the aortic root diameter after the withdrawal of ALO following PA1 and PE treatments (PA1<ALO and PE<ALO, respectively; Fig. S1). When 6-month-old MFS animals were subjected to PA1 treatment with ALO and subsequently withdrawn from the drug for three months, until reaching 9 months of age, no statistical difference was obtained between their respective aortic root diameters (dashed red boxes of PA1<ALO 9-month-old MFS groups in Fig. S4A; Table S6). On the other hand, the 3-month-old animals subjected to PE treatment with ALO were withdrawn from the drug until reaching 6 months of age (WT and MFS groups labeled with red circles in Fig. S4B). There was no statistical difference between ALO-treated MFS groups regardless of whether ALO was subsequently withdrawn or not (p=0.08) (dashed brown boxes of PE<ALO 6-month-old MFS groups in Fig. S4B; Table S7). Therefore, results show that the inhibitory effect of ALO on the aneurysm is only effective while the drug is administered.

### Allopurinol prevents endothelial dysfunction in MFS ascending aorta

ROS generated by endothelial XOR could mediate alterations in the endothelial-associated vascular function by reducing NO bioavailability via direct interaction with O_2_^-^^32, 33, 50^. Previous evidence indicated that the MFS aorta shows endothelial dysfunction^14, 51, 52^. Thus, we analyzed the potential therapeutic effects of ALO treatment on endothelial function (Fig. 4). To this aim, we used wire-myography to measure the reactivity of endothelium-intact ascending aortas from 9-month-old MFS mice (PA2; Fig. S1), age at which, in our hands, endothelial dysfunctions were more clearly observed^52^. Firstly, we examined aortic root diameter in this mouse subset and the aneurysm was also inhibited after seven months of ALO treatment (PA2) (Fig. S5 and Table S8). Thereafter, ascending aortae were isolated, vessels were settled in the myograph and contracted with KCl to check their functionality (Fig. 4A). All aortae responded similarly to KCl (Fig. 4B and Table S10). The relaxant response to acetylcholine (ACh; Fig. 4A), mostly mediated by activation of endothelial NOS (NOS3), is an indicator of endothelial function. Untreated MFS mice showed a reduced sensitivity (pEC_50_) to endothelium-dependent ACh-induced relaxation compared with WT (the curve shifted to the right in Fig. 4C/asterisks; Table S9), which indicated that ACh-mediated aortic relaxation was impaired in MFS mice and, therefore, demonstrative of dysfunctional endothelium. The treatment with ALO showed an ACh-induced relaxation in MFS aortae that was indistinguishable from WT animals (pEC_50_ 7.49 ± 0.19 and 7.52 ± 0.10, respectively) (Fig. 4C; Table S9), proving that ALO treatment normalizes endothelial function in MFS mice.

**Figure 4.**
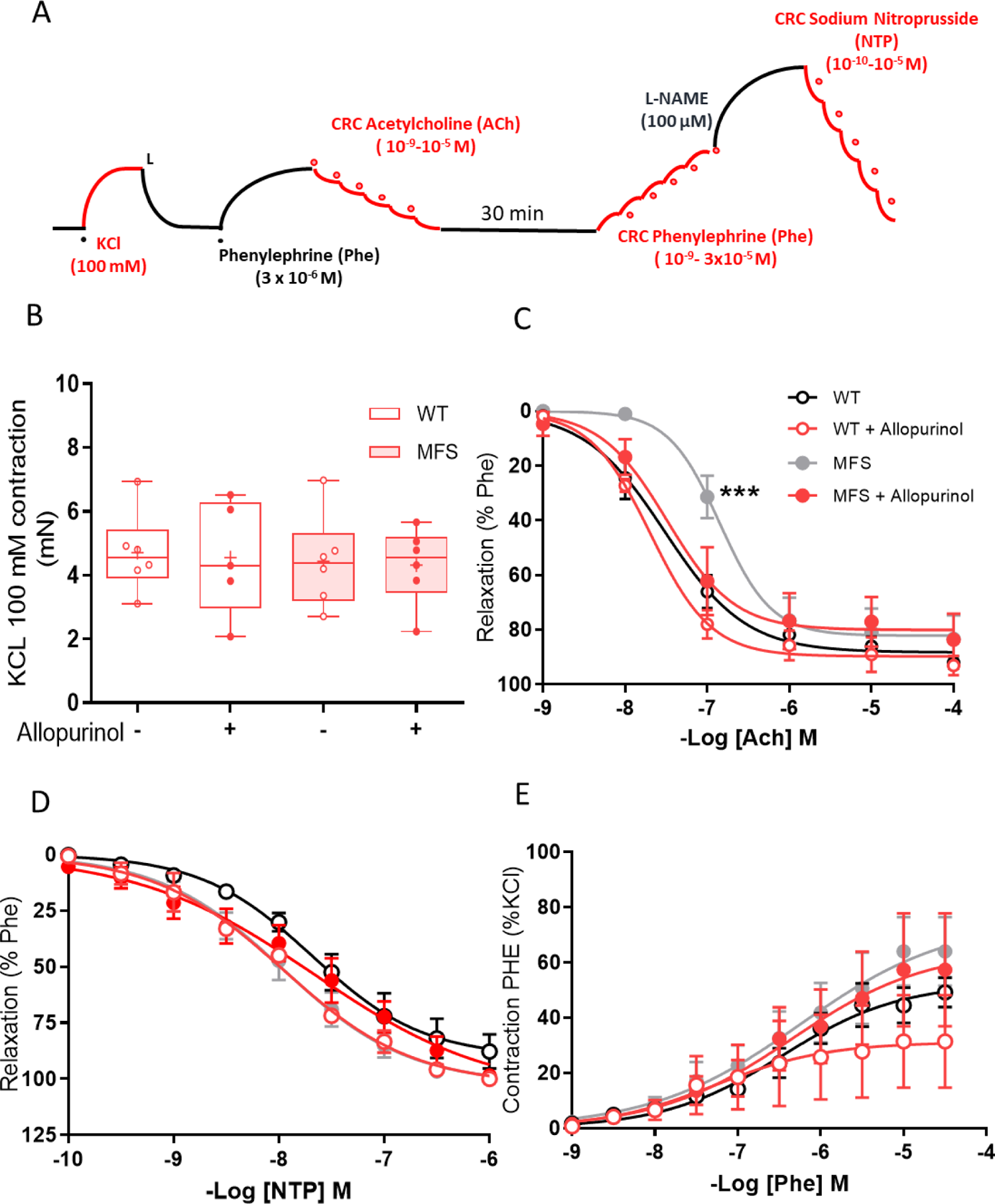
Allopurinol prevents the endothelial vasodilation dysfunction of MFS aorta. **(A)** Experimental protocol for wire-myography experiments carried out in isolated 9-month-old ascending aorta from the diverse experimental groups indicated in the following panels. Red lines indicate where myographic measurements were obtained. **(B)** Physiological state test of aortic reactivity following the addition of KCl (100 mM). Data represented as boxplots. See also Table S10. **(C)** Relaxation of aortic reactivity following ACh addition. **(D)** Relaxation of reactivity after the addition of the NO donor sodium nitroprusside. **(E)** Contraction of aortic reactivity after the addition of phenylephrine. Data in C-E are mean ± SD. See also Tables S9 and S10. Statistical test tests: Two-way ANOVA and Tukey post-test (B); non-linear regression of data by fitting Phe/ACh/NTP concentration-response curves to a sigmoidal dose-response (C-E). \*\*\**P*≤0.001.

To check the participation of the NO/GC/cGMP pathway in the VSMCs of untreated MFS mice, we next added increased concentrations of the NO donor sodium nitroprusside (NTP) to aortae pre-contracted with phenylephrine (Phe) and the non-selective NOS inhibitor N(gamma)-nitro-L-arginine methyl ester (L-NAME; Fig. 4A), resulting in similar concentration-dependent aortic vasodilatations in untreated MFS and WT mice (Fig. 4D). No changes in the vasorelaxant response to NTP were observed following ALO administration either in WT or MFS mice (Fig. 4D; Table S9). We also evaluated the α_1_-associated contractile response of ascending aortae when stimulated with different concentrations of Phe. Results were highly similar between non-treated MFS and WT ascending aortae or following ALO treatment (Fig. 4E; Table S10).

At this point of the study, there is an apparent inconsistency knowing that allopurinol is greatly inhibiting aortic aneurysm over the age in MFS mice (examined at 3-to-9 months old animals), but nevertheless XOR is only significantly increased in young animals (3 months-age). This means that other mechanisms directly or indirectly mediated by allopurinol must be affected. This conundrum is resolved below.

### Allopurinol prevents increased levels of H_2_O_2_ in MFS aorta

Considering that ALO is clinically prescribed to treat hyperuricemia, we first measured UA and allantoin levels in blood plasma in WT and MFS mice to see potential changes associated with genotype or age or both. We noticed that neither UA nor allantoin, nor their ratio, showed any change between WT and MFS mice over age (Fig. S6A-C). Next, we analyzed whether ALO (PA1) reduced UA in blood plasma, but we observed that UA levels remained unaltered (Fig. S7). The possibility that the inhibition of aneurysm progression was mediated by reducing blood pressure as previously reported in other studies^53^ was also discarded since systolic blood pressure values also remained unaltered (Fig. S8).

The aortopathy inhibition by ALO inhibiting XOR activity only happens in the early ages of MFS mice (3 months), but it cannot be explained in older animals. Therefore, another mechanism(s) must significantly participate in which redox stress should be directly or indirectly affected by ALO. In this regard, it was reported that ALO could directly scavenge ROS^54^, thus, we measured ROS levels in blood plasma and aorta from WT and MFS mice. We chose to measure H_2_O_2_ because it is the major ROS produced both by XOR under aerobic conditions^55^, which occur in the heart and aorta, and by NOX4, which is upregulated in MFS aorta^17^. In blood plasma, we observed no differences with age between WT and MFS mice (Fig. S9). In contrast, ascending aorta rings from MFS mice showed significantly higher levels of H_2_O_2_ compared with WT mice of different ages (Fig. 5A). The *in vitro* administration of ALO to these aortic rings normalized H_2_O_2_ levels. This was also the case when ALO was administered *in vivo* in MFS mice (PE), whose aortic H_2_O_2_ levels were highly similar to those of untreated WT aortae (Fig. 5B). Therefore, ALO either prevented or inhibited the MFS-associated high levels of H_2_O_2_ in the ascending aorta irrespectively of *in vivo* or *in vitro* administration.

**Figure 5.**
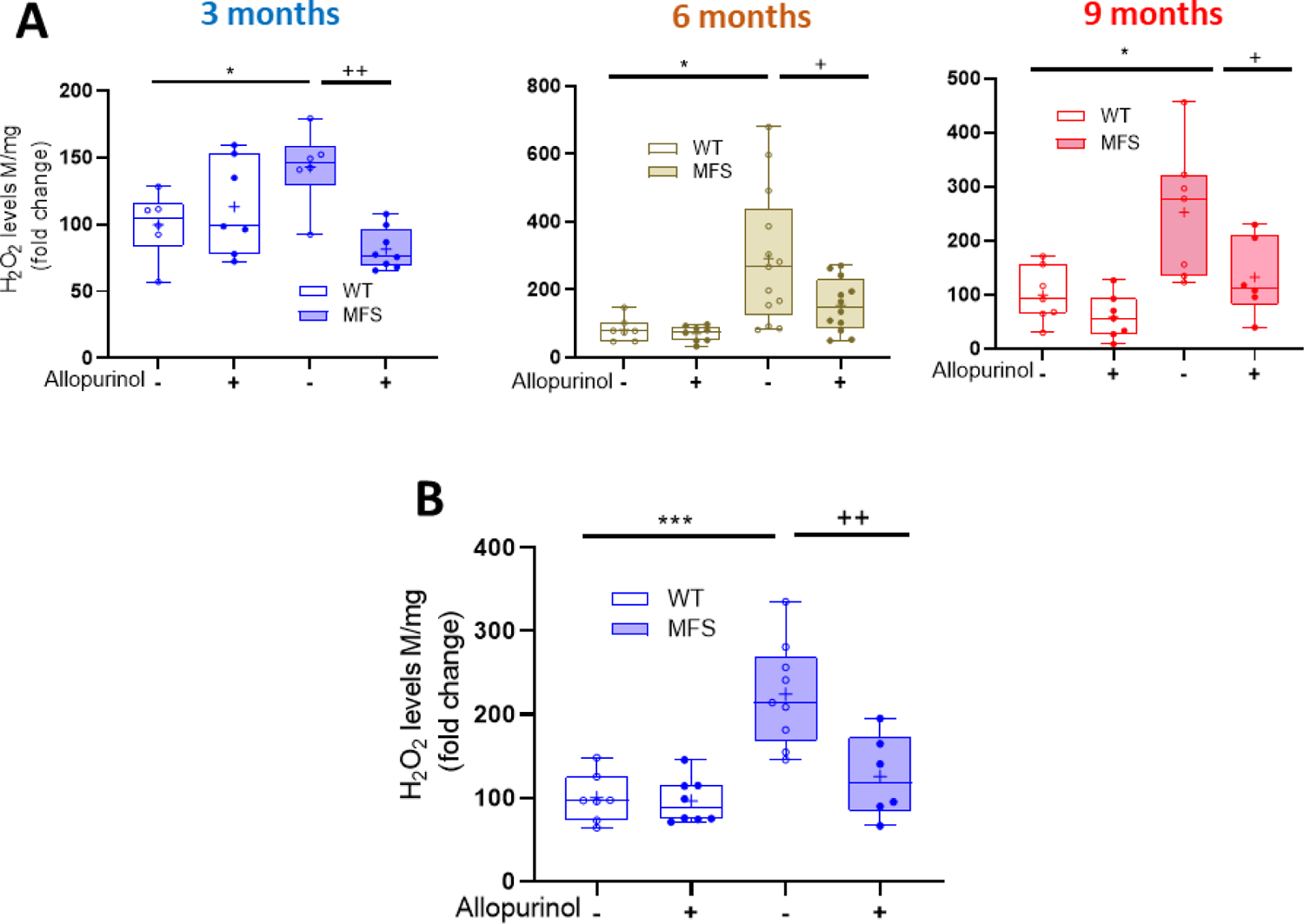
High H_2_O_2_ levels generated in the ascending aorta of MFS mice are inhibited by allopurinol when administered both *in vitro* and *in vivo*. (**A**) H_2_O_2_ levels produced in ascending aortic rings over age in WT and MFS mice. Allopurinol was added *in vitro* to the assay containing aortic rings and the fluorescence signal measured after 2 h. (**B**) H_2_O_2_ levels measured in ascending aortic rings of WT and MFS mice preventively (PE) treated with allopurinol (*in vivo*). Data represented as boxplots. Statistical test analysis: Two-way ANOVA and Tukey’s post-test \*\*\**P*≤0.001, ^++^*P*≤0.01, *^/+^*P*≤0.05; *effect of genotype; ^++^effect of treatment.

### Allopurinol downregulates NOX4 and MMP2 expression in MFS aorta

Knowing that ALO blocks aortic aneurysm progression with age (3-, 6- and 9-month-old MFS mice), that aortic H_2_O_2_ levels also remain significantly elevated at these ages, and that XOR is only upregulated at early ages (3 months), we postulated that, regardless of its scavenging action, ALO could be directly or indirectly affecting other enzymatic H_2_O_2_ sources at the MFS aorta. In this regard, we next examined whether ALO treatments affected NOX4 expression, which greatly and directly produces H_2_O_2_ and whose expression and activity are abnormally increased in MFS aortae^17, 18^ Thus, we evaluated NOX4 mRNA levels in the aorta of WT and MFS mice palliatively (PA1) and preventively (PE) treated with ALO (Figs. 6). In both cases, ALO-treated MFS mice inhibited the characteristic upregulation of NOX4 in MFS aorta (Fig. 6A and B).

**Figure 6.**
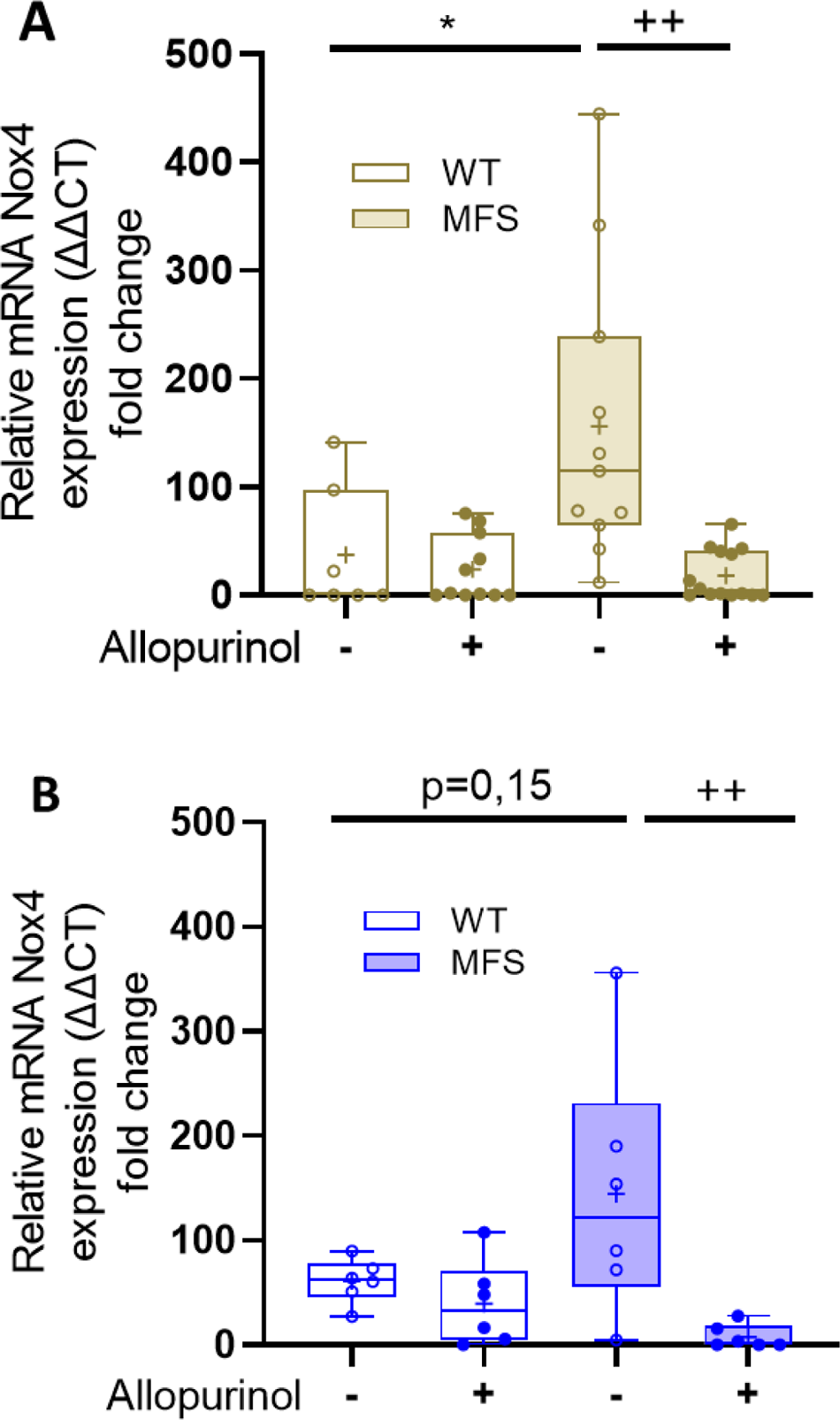
Allopurinol prevents the upregulation of NADPH oxidase NOX4 in MFS aorta. mRNA expression levels of NOX4 in WT and MFS mice following the PA1 **(A)** or PE **(B)** allopurinol treatments. N=7-12. Statistical test analysis: Kruskal-Wallis and Dunn’s multiple comparison test. *^/+^*P*≤0.05 and ^++^*P*≤0.01; *effect of genotype; ^+^effect of treatment.

Another potential and complementary mechanism by which ALO, acting as an indirect antioxidant, could contribute to explaining the large reduction of MFS-associated aortic wall disarrays is by inhibiting the ROS-mediated activation of MPPs, as ROS are potent activators of MMPs^56^. MMP2 and MMP9, two well-known MMPs involved in MFS aortopathy^57^ can be indirectly activated by ROS from their inactive-zymogen isoforms and via direct activation of their gene expression^58^. As expected, MFS aortae showed increased MMP2 protein levels in the tunica media compared with WT aorta. Both PE and PA1 ALO treatments inhibited the increased expression of MMP2 (Fig. 7).

**Figure 7.**
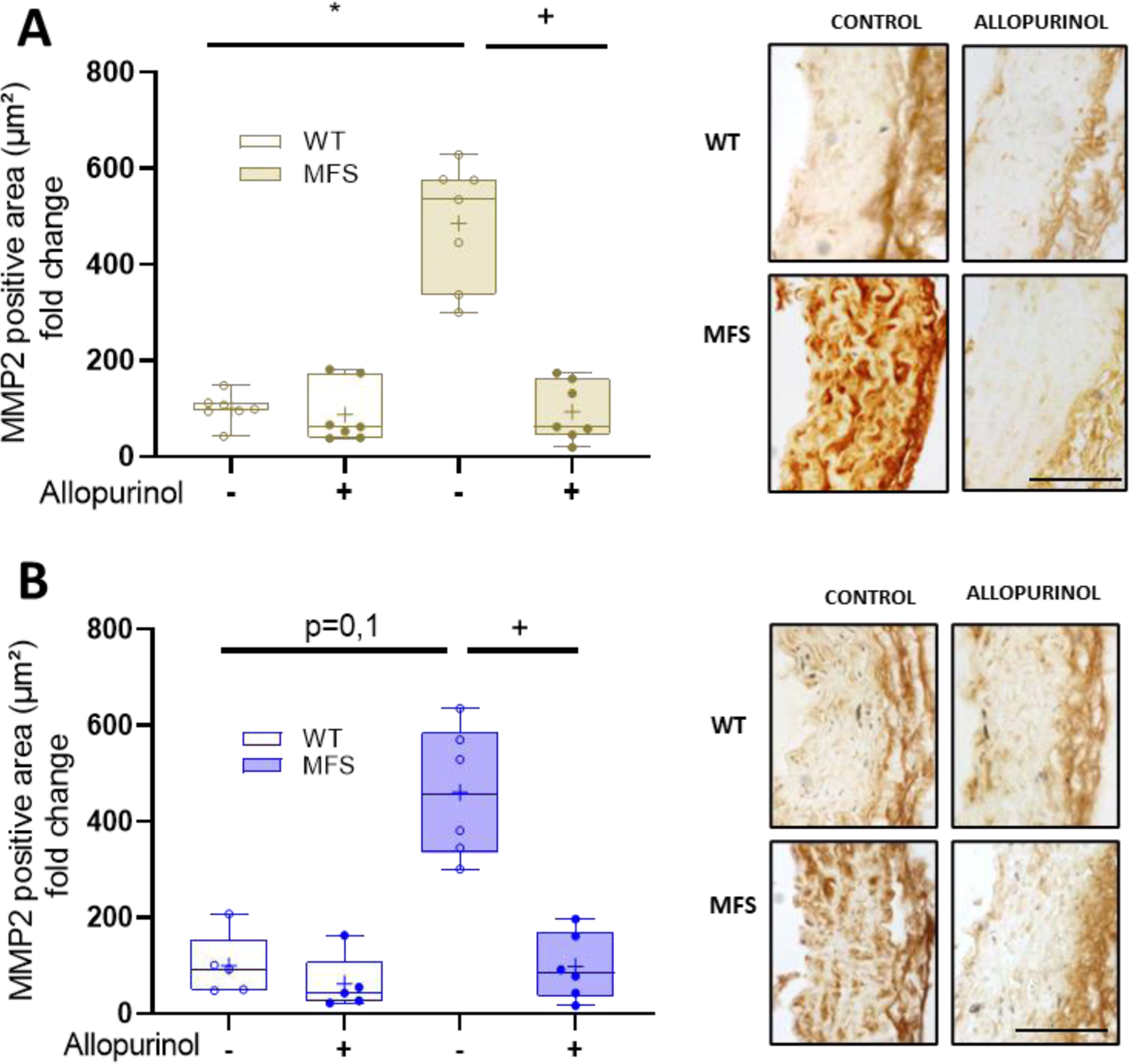
Allopurinol reduces increased MMP2 levels in the tunica media of MFS aorta. MMP2 levels in the tunica media revealed by immunohistochemistry with anti-MMP2 antibodies in paraffin-embedded aortae from WT and MFS mice following palliative (PA1 **(A)** or preventive (PE**) (B)** administration of allopurinol. N= 6-8. On the right, quantitative analysis of the respective HRP immunostaining. Bar, 100 µm. Statistical test analysis: Two-way ANOVA and Tukey’s post-test *^/+^*P*≤0.05. *Effect of genotype; ^++^effect of treatment.

### Allopurinol prevents redox stress-associated injuries in the aortic tunica media of MFS mice

RNS can be produced due to the interaction of endothelial-generated NO and XDH-induced ROS^32, 33^. Peroxynitrite and 3’-nitrotyrosine (3-NT) products are reliable redox stress biomarkers; immunohistochemical evaluation showed greater levels of 3-NT in the tunica media of MFS aortae. In both PA1 and PE approaches, ALO significantly prevented this increase (Fig. S10A and B).

The nuclear factor erythroid 2-related factor 2 (NRF2) is a key transcription factor that regulates the expression of several antioxidant defense mechanisms. Oxidative stress triggers its phosphorylation (pNRF2), being subsequently translocated to the nucleus to activate the expression response of physiological antioxidant enzymes^59^. Thus, we evaluated the nuclear presence of pNRF2 in aortic paraffin sections from WT and MFS mice treated with ALO after PA1 and PE treatments (Fig. 8A and B, respectively). Aortic media showed a higher presence of nuclear pNRF2 in MFS than WT VSMCs. This result demonstrated that the MFS aorta suffered redox stress, which in turn triggered the endogenous antioxidant response. However, when ALO was administered to MFS mice, the number of pNRF2 stained nuclei was indistinguishable from WT aortae, regardless of the experimental treatment followed. This result is a logical consequence of the ALO-induced inhibition of the oxidative stress occurring in the MFS aorta.

**Figure 8.**
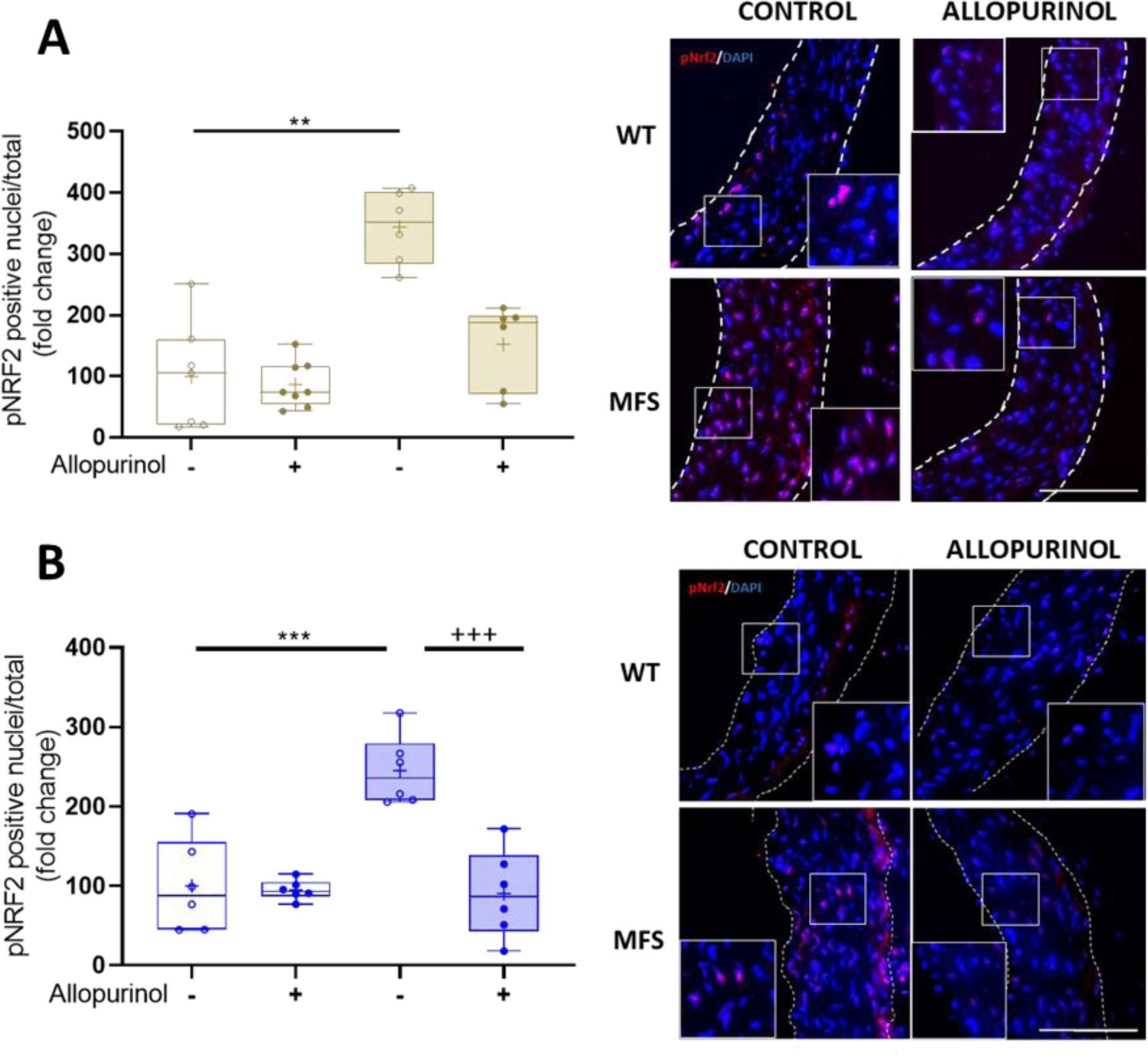
Allopurinol inhibits the oxidative stress-associated increased nuclear translocation of pNRF2 in MFS aorta. Quantitative analysis and representative images of the nuclear translocation of the phosphorylated form of NRF2 in VSMCs of the tunica media of WT and MFS mice treated with allopurinol palliatively (PA1) **(A)** and preventively (PE) **(B)**. Insets in the immunofluorescent images detail the nuclear localization of pNRF2. Bar, 100 µm. Statistical test: Kruskal-Wallis and Dunn’s multiple comparison tests. ***^/+++^*P*≤0.001 \*\**P*≤0.01; *effect of genotype; ^+^effect of treatment.

Elastic fiber disruption in the MFS aorta is often accompanied by compensatory fibrotic remodeling supported by collagen overexpression and/or organization rearrangements occurring mainly in the tunica media^60^. This remodeling can be visualized under a polarized light microscope in paraffin-embedded aortae cross-sections stained with Picrosirius red^61^. Green and red collagen fibers were observed, which is illustrative of their different fiber thickness and assembly compaction and, therefore, of their degrees of maturity^62^. We observed an increase in collagen fibers in the aortic tunica media of 6-month-old MFS mice (PA1 treatment), mainly due to a significant increase in immature (green) fibers. This change is demonstrative of a fibrotic-like response. Notably, this collagen-associated reorganization was not produced after ALO treatments (PA1; Fig. 9). No differences were achieved in preventive ALO administration in MFS mice even though similar changes in green fibers tend to occur as well (Fig. S11). In PA1 and PE treatments, ALO did not reduce the increased trend of mature (red) collagen fiber formation (Fig. 9 and Fig. S11, respectively). These results indicated that ALO also reduced the characteristic fibrotic response taking place in the aortic media of MFS mice.

**Figure 9.**
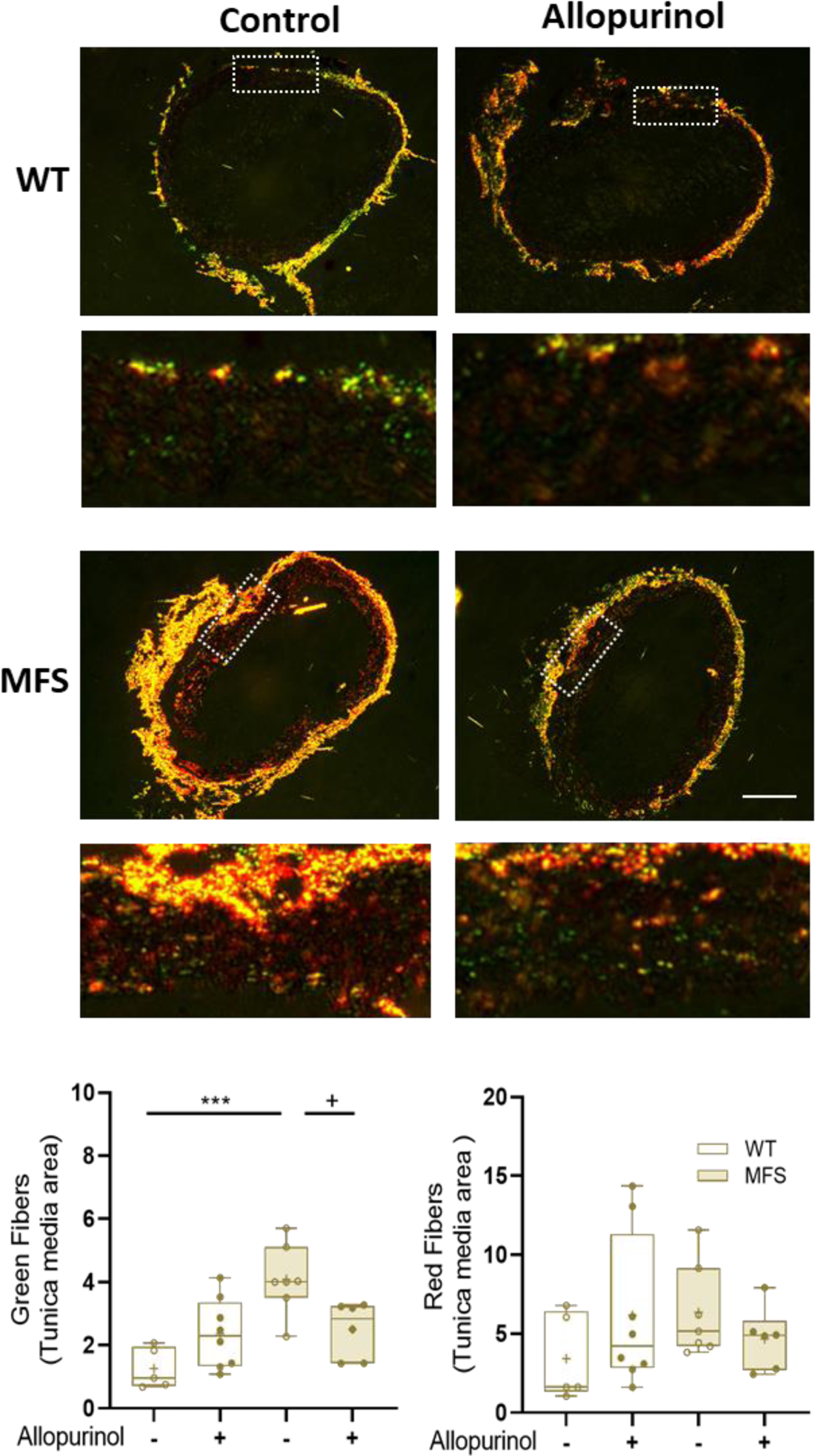
Allopurinol reduces the MFS-associated fibrotic remodeling of the tunica media. Immature (green) and mature (red) collagen fibers of the tunicae media and adventitia of WT and MFS aortae stained with Picrosirius red visualized under the polarized microscope. WT and MFS mice were palliatively treated (PA1) or not with allopurinol (n=5-8). Representative fluorescent images of the whole aorta and enhanced indicated regions (white dashed lines). In enhanced images, the adventitia in located above and the media just below. Bar, 100 µm. The respective quantitative analysis of both types of collagen fibers is shown under the images. Statistical test: Kruskal-Wallis and Dunn’s multiple comparison tests. \*\*\**P*≤0.001 and ^+^*P*≤0.05; *effect of genotype; ^+^effect of treatment.

## DISCUSSION

Despite the significant insights reported over the last two decades regarding the molecular mechanisms that participate in the formation and progression of aortic aneurysms in MFS, current pharmacological approaches (losartan, atenolol, or both together) have proven insufficient to halt or mitigate aortic aneurysm progression^63^. Keeping this in mind, the aim of our study was, on the one hand, to obtain further insights into the contribution of redox stress to the molecular pathogenesis of aortic aneurysms, focusing on XOR, which, jointly with NADPH oxidases, is the most potent ROS-generating system in the cardiovascular system in health and disease; on the other hand, we aimed to obtain solid experimental evidence in a representative murine model of MFS to support a new pharmacological approach targeting XOR-associated oxidative stress (allopurinol), which could be translated to MFS patients.

This article produced several main findings: (i) XOR is upregulated in the aorta of both MFS patients and mice; (ii) this upregulation is accompanied by increased enzymatic activity of the oxidase (XO) form in detriment of the dehydrogenase (XDH) one; (iii) the XOR inhibitor allopurinol (ALO) prevents both the formation and progression of aortic root dilation in MFS mice; (iv) this inhibitory effect is non-permanent since the withdrawal of ALO causes the reappearance of the aneurysm; (v) ALO prevents the characteristic MFS endothelial-dependent vasodilator dysfunction; (vi) ALO inhibits the MFS-associated increase of aortic H_2_O_2_ levels both *in vivo* and *in vitro* as well as the subsequent accompanying redox stress-associated reactions, such as accumulation of 3-NT, augmented nuclear translocation of pNRF2 and collagen-associated fibrotic remodeling of the aortic media; and (vii) ALO inhibits MFS-associated upregulation of NOX4 and reduces MMP2 expression in the aortic tunica media. Mechanistically, we postulated that ALO mitigates MFS aortopathy progression by acting as an antioxidant both directly scavenging ROS and inhibiting XOR activity, and indirectly downregulating NOX4 and MMP2 expression.

In aortic samples from MFS patients, XOR expression increased compared with healthy controls. This result was confirmed in MFS mice both at the protein and mRNA level. These changes were only observed in young (3-month-old) but not in older mice (6- and 9-month-old). It is possible that XOR upregulation only occurs while the aortic dilation undergoes rapid growth, which we found to occur until 3 months of age, becoming slower in older animals^45^. Nonetheless, no differences between WT and MFS aortae were observed regarding XOR activity in the study of the reactivity of MFS aorta^14^, and after nitro-oleic acid treatment as a mediator in ERK1/2 and NOS2 expression in MFS aortopathy^64^. Explanations for this apparent discrepancy may be related to the different murine MFS models used (mgR/mgR and AngII infusion-induced acceleration of the aortopathy in MFS mouse, respectively) or the different XOR activity assays utilized, and/or even to local animal facility conditions.

ALO is a well-characterized specific inhibitor of XOR activity, mainly of the XO form^42^, which was confirmed in the XOR enzymatic assay (Fig. S2). The main clinical actions of ALO are widely linked to UA-associated pathologies^65, 66^. UA plasma levels were highly similar between MFS and WT animals, which at first glance discarded a UA-mediated effect in the aortopathy in MFS mice. Curiously, ALO did not cause the expected reduction of UA and/or allantoin plasma levels. However, it is important to highlight that the equivalent dose of ALO used here in mice (20 mg/Kg/day) if administered to humans (∼120 mg/day calculated for a weight of 70 Kg)^67^ does not reduce normal plasma UA levels (≤6-7 mg/dL)^68^. Therefore, the blockade of MFS aortopathy by ALO is, in principle, not directly related to changes in UA, but to another accompanying mechanism(s).

Of note, ALO has a ROS lowering effect not necessarily related to its XOR inhibition activity, acting accordingly as a direct ROS scavenging moiety^68–71^. We demonstrated this possibility when ALO was added *in vitro* to MFS aortic rings, quickly attenuating their intrinsic increased H_2_O_2_ production levels, regardless of mouse age. Importantly, this reduction was also observed in aortic rings from MFS mice with the drug administered *in vivo*. In addition to the direct scavenging role of ALO, its effectiveness as a direct antioxidant can be reinforced, reducing the characteristic aortic upregulation of NOX4, knowing that NOX4 is the NADPH oxidase that directly generates H_2_O_2_^72^. Pathological processes that usually accompany redox stress, such as the formation of 3-NT, pNRF2 nuclear translocation and fibrotic remodeling responses were all reduced or inhibited by ALO. The pathophysiological significance of these results in the MFS aortic media suggested several conclusions: (i) the accumulation of 3-NT is representative of abnormal RNS formation because of the pathological uncoupling of NO, which in turn leads to protein nitration via the formation of the highly reactive intermediate peroxynitrite and its subsequent product 3-NT^73^. This 3-NT upsurge in MFS aortic media is consistent with the recent demonstration of NO uncoupling in aneurysm formation in both MFS mice and patients^64, 74, 75^. Along this line of evidence, we identified actin as a nitrated protein target in MFS aorta from MFS patients^17^, contributing, in this manner, to the reported damaged contractile properties of VSMCs in MFS^76^; (ii) redox stress is so elevated in MFS aorta that the intrinsic physiological antioxidant response mediated by the nuclear translocation of pNRF2^60^ is not high enough to compensate it. In contrast, ALO treatment allows this compensation, normalizing, to a large extent, the endogenous redox levels in MFS aorta and associated modifications; and (iii) ALO normalized, to a great extent, the content of immature collagen (green) fibers, hence mitigating the fibrotic remodeling response that usually accompanies the aneurysm to compensate elastic fiber disarray. It would be interesting to know whether ALO is also able to prevent the characteristic phenotypic switch of VSMCs to a mixed contractile-secretory phenotype.^77, 78^

Of notice, ALO normalized ACh-stimulated vasodilator function, improving endothelial function. It is not a surprise given the reported effects of ALO on increasing NO bioavailability, totally or partially attributable to a reduction in ROS production.^32, 33^

In reference to the normalization by ALO treatments (PA1 and PE) of the structural aortic wall organization evidenced by the normalization of elastic laminae breaks, this effect could be explained by the inhibition of the ROS-mediated activation of MMP2 upregulation, which, together with MMP9, are two well-known MMPs upregulated and overactivated in MFS aorta^58^. Nonetheless, XOR itself can directly activate MMPs in a ROS-independent manner^79^.

Finally, it cannot be discarded that ALO could also mediate its inhibitory action on aortopathy by acting as an anti-inflammatory agent^80^ and/or also indirectly blocking the ROS-induced heat shock protein expression generating endoplasmic reticulum (ER) stress, which has an impact on abdominal and thoracic aortic aneurysms^81–84^. These two possibilities deserve future attention.

We are aware that our study has some limitations: (i) ALO has a short half-life in plasma and is rapidly metabolized to oxypurinol, which has a longer lifespan^42^. It is possible that oxypurinol rather than allopurinol might be the primary metabolite that directly mediates the results we report herein. Nonetheless, in this respect, it is important to take into account that it is not recommended to administer higher doses of ALO in humans (>300 mg/day) because the high concentration of oxypurinol generated then acts as a pro-oxidant instead of antioxidant, favoring oxidative stress^68^; (ii) the MFS model used in our study (C1041G) is a very useful model to evaluate the temporal course of aortopathy (besides other clinical manifestations), but it is not the most suitable to evaluate the survival rate after ALO treatments. Other, more severe MFS murine models for aortic dissection and rupture, such as the *Fbn1* hypomorphic mouse (mgR/mgR)^85^ or the AngiotensinII-induced accelerated model of MFS aortopathy^64^, both leading to aortic dissection and rupture, would be more appropriate experimental murine models for this aim; and (iii) the aortic arch and descending thoracic aorta have not been studied; (iv) we have only studied ALO’s effects on the cardiovascular system (TAA more specifically) because it is responsible for the life-threatening complication in MFS. Nevertheless, since MFS is a multisystemic disorder, it would also be of interest to study the putative effects of ALO on the other systems affected, even though it has been proven to be a safe and well-tolerated drug.

## CONCLUSION

Our results definitively place redox stress among the molecular mechanisms that actively participate in the pathogenesis of aortic aneurysm in MFS by the persistent high production levels of ROS mediated by upregulation of XOR (current study), the NADPH oxidase NOX4^17^, eNOS dysfunction^74^ and mitochondrial electron transport-associated redox stress^86^, all from studies performed both in MFS murine models and patients. How, and to what extent, do each of these interlinked mechanisms participate in aortic injury is currently unknown, but most likely they all act additively or synergistically to damage the aorta severely and permanently. We here demonstrated ALO’s effectiveness, acting as a strong antioxidant drug in MFS aortae directly both as an inhibitor of XOR activity in early ages when XOR is overexpressed, and as ROS scavenger in the absence of XOR activity, and indirectly by downregulating NOX4 and MMP2 overexpression. ALO is an economic drug, which has been widely prescribed in clinical practice since the latter half of the last century and, most importantly, has been proven to be effective, safe, and usually well tolerated. ALO has been reported to present lower rates of cardiovascular events (related to hypertension) compared with non-treated patients^87^. Of note, a preprint study reported that febuxostat has similar effects to ALO, blocking MFS aortopathy.^88^ Febuxostat is another specific XOR inhibitor not resembling purines or pyrimidines^89^. Some concerns regarding cardiovascular safety have been reported for febuxostat compared with ALO^90^, but a recent study seems to refute this concern^91^. In sum, our results in MFS mice support the design of a clinical trial with ALO as having a potential beneficial effect in the pharmacological clinical practice of MFS patients. Last, but not least, such a potential benefit could only be obtained when the drug is administered chronically.

## Acknowledgments

G.E. dedicates this article to the memory of Professor Mercedes Durfort i Coll (1943-2022). We deeply thank Isabel Fabregat and Hal Dietz for reviews and comments on previous versions of the manuscript, Ana Paula Dantas (IDIBAPS) and Coral Sanfeliu (CSIC) for helpful methodological advice with H_2_O_2_ fluorometric measurements, Helena Kruyer for patient editorial assistance, and Maria Encarnación Palomo and María Teresa Muñoz for excellent technical assistance and lab management, respectively.

## Author contributions

Conceptualization: GE; Methodology: IR-R, CA, KdR, BP, AC, MA, FJ-A. Investigation: IR-R, KdR, CA, BP, MA, VC, FJ-A, GE; Funding acquisition: GE, FJ-A and MCG-C; Supervision: GT-T, MCG-C, VC, FJ-A, GE; Writing – original draft: GE; Writing – review & editing: all authors.

## Nonstandard abbreviations

ALO: allopurinol

CRCs: cumulative concentration-response curves

3-NT: 3’-nitrotyrosine

PA: palliative experimental treatment with allopurinol

PE: preventive experimental treatment with allopurinol

PE<ALO and PA<ALO: respective PE and PA treatments with subsequent allopurinol withdrawal.

## SUPPLEMENTARY MATERIAL

### Supplemental Methods

### Supplemental Figures

**Figure S1.**
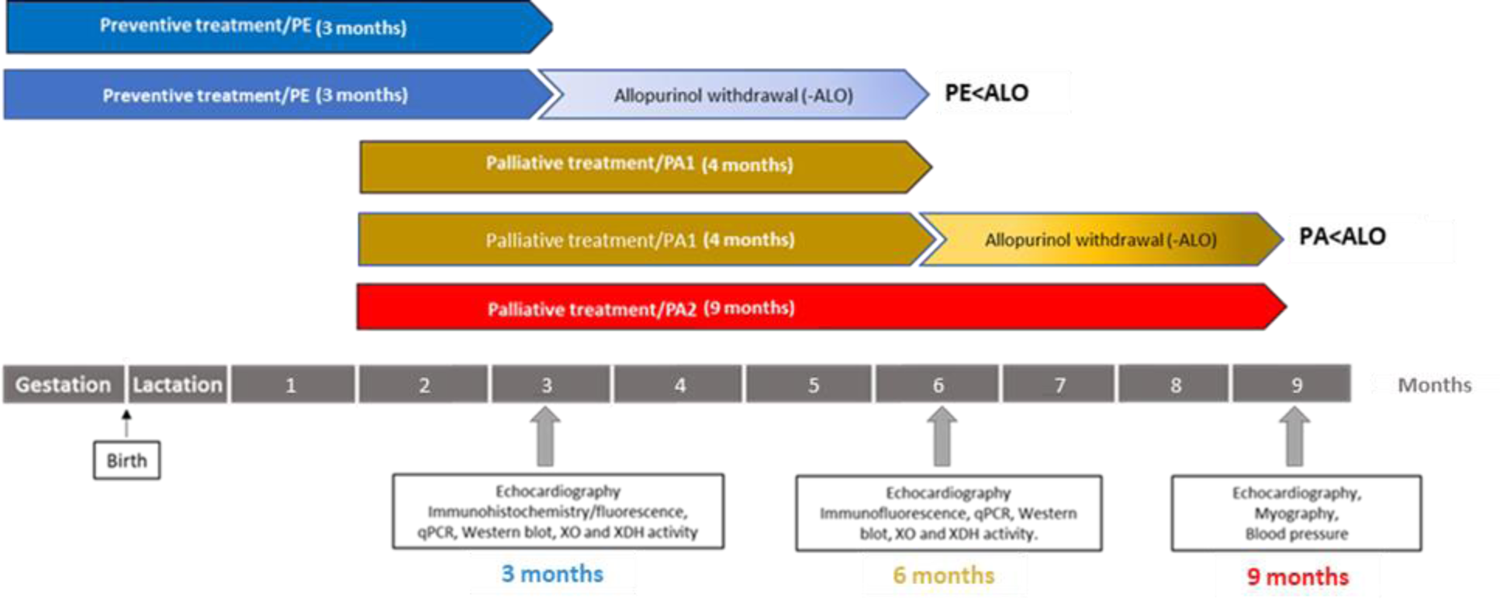
Representative scheme of the experimental protocols for allopurinol treatments. PE: preventive allopurinol (ALO) treatment (endpoint at 3-month-old); PA: palliative allopurinol treatments PA1 and PA2 whose difference between them being the duration of treatment, which was 4 months (endpoint at 6-month-old mice) and 7 months (endpoint at 9-month-old mice), respectively. PE<ALO: preventive allopurinol treatment followed by the subsequent drug withdrawal for 3 months (endpoint at 6-months-old); PA1<ALO: palliative allopurinol treatment followed by the subsequent drug withdrawal for 3 months (endpoint at 9-month-old). Note that each endpoint duration protocol has a different code color, which is maintained for the figures and supplemental figures shown throughout the manuscript: blue (PE), brown (PA1) and red (PA2).

**Figure S2.**
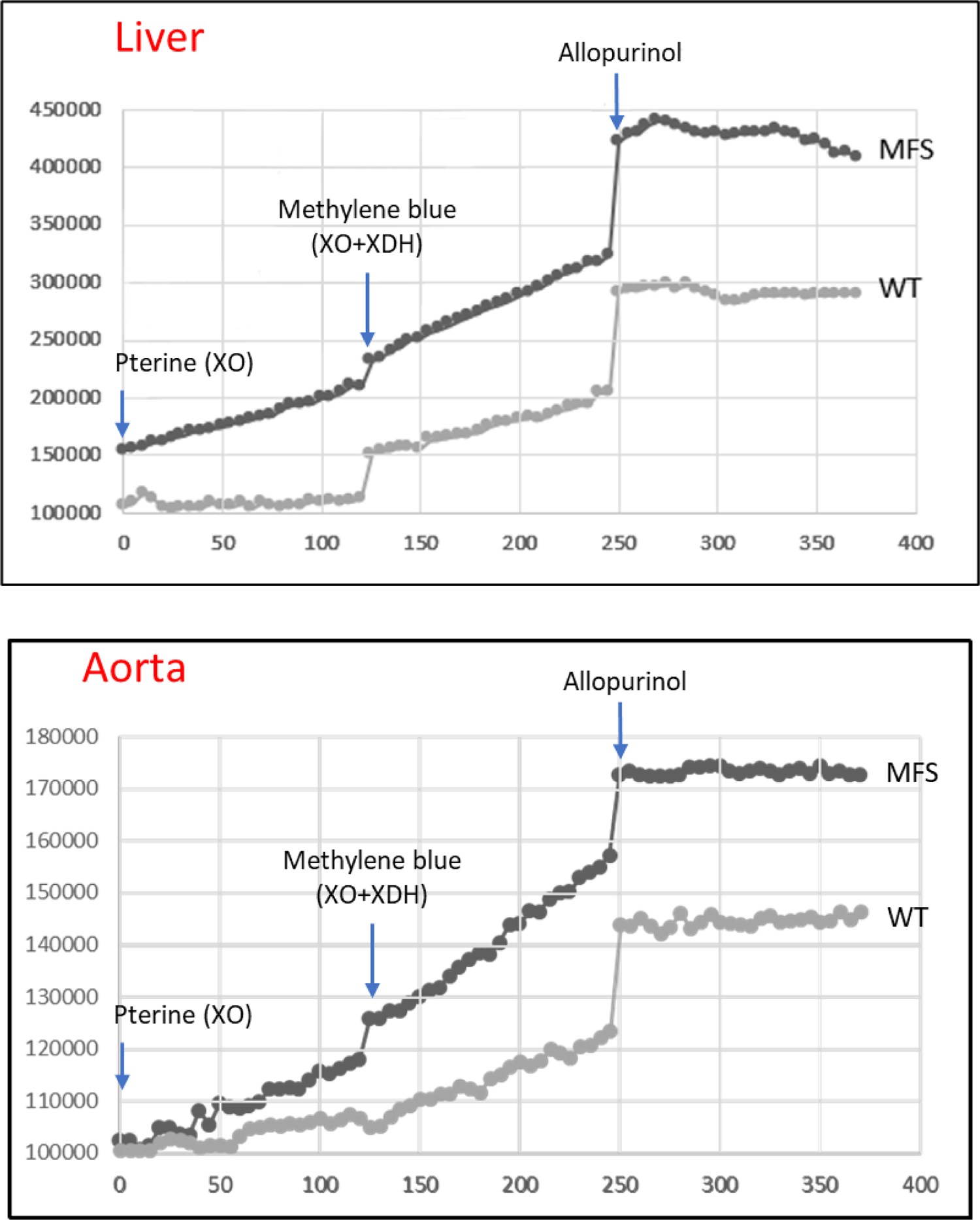
Fluorometric assay for measuring xanthine dehydrogenase (XDH) and oxidase (XO) enzymatic activity in liver and aortic tissues. Liver (as positive control of the assay) and aortic WT and MFS lysates had pterine added as a specific substrate for XO and methylene blue for total XOR (XO+XDH) activity. Allopurinol was added at the end of the assay to check its effectiveness in inhibiting total XDH activity. Units in the Y axis are fluorescence arbitrary units.

**Figure S3.**
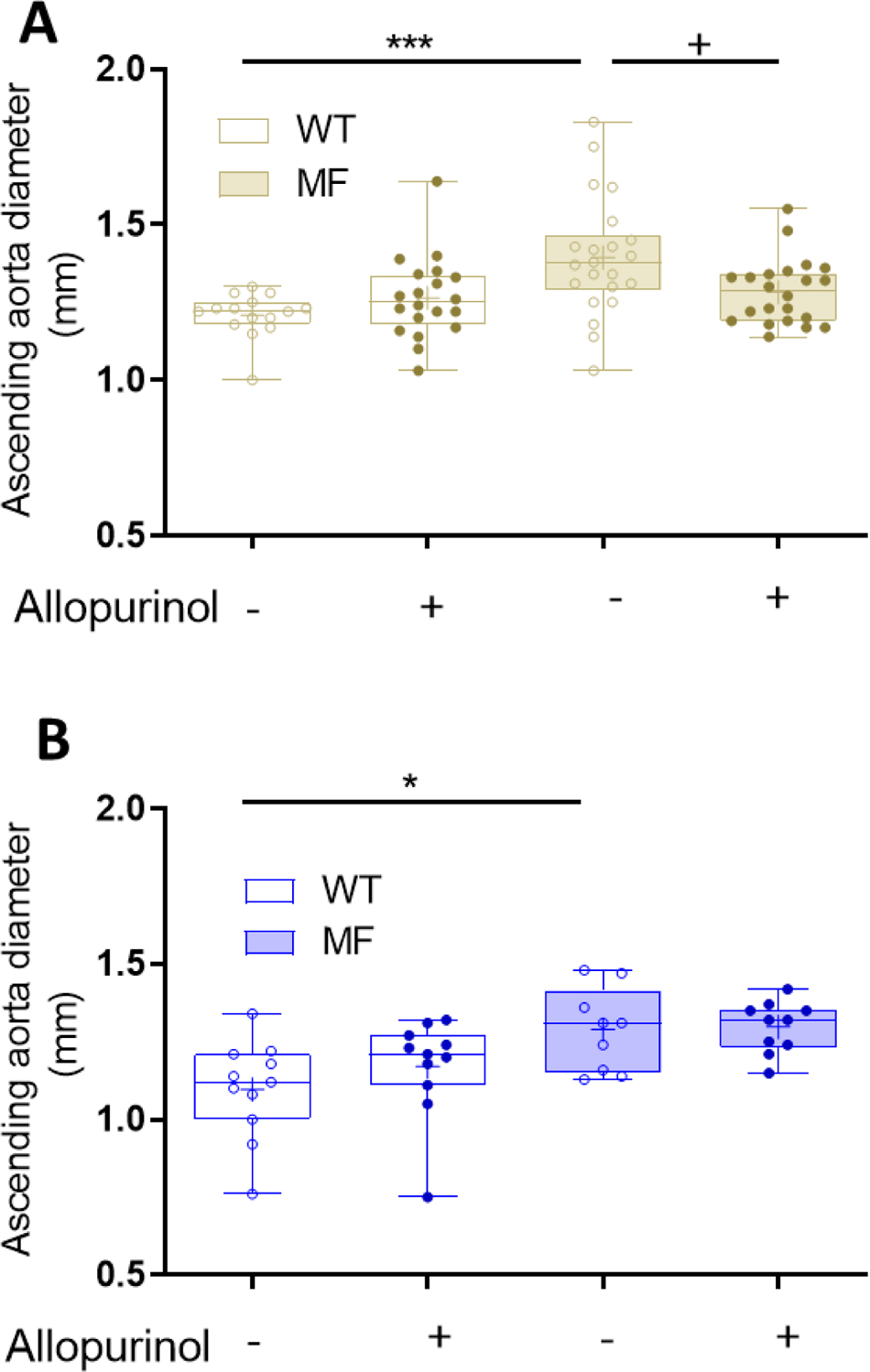
Ascending aorta diameter in WT and MFS mice treated with allopurinol. **(A)** Ultrasonography in WT and MFS mice of 6 months of age palliatively treated with allopurinol (PA1). See also Table S4. (**B)** Ultrasonography in 3 months of age WT and MFS mice preventively treated with allopurinol (PE). See also Table S5. Data represented as boxplots. Statistical analysis: Two-way ANOVA and Tukey post-test (A and B). \*\*\**P*≤0.001 and \**P*≤0.05; *effect of genotype; ^+^effect of treatment.

**Figure S4.**
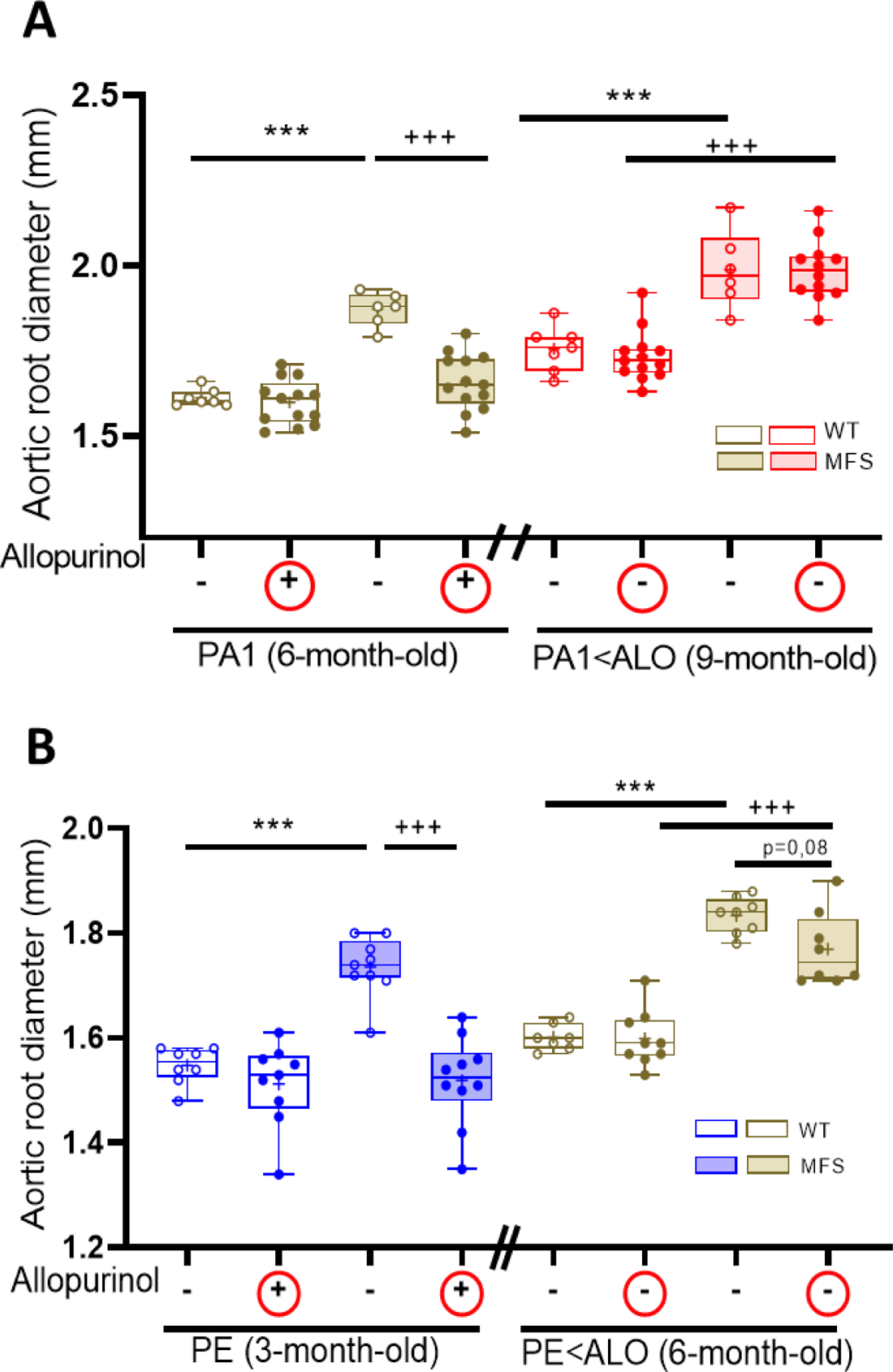
Reversion of aortic root dilation following the withdrawal of allopurinol. (**A**) WT and MFS mice were treated with allopurinol (ALO) following the palliative treatment from 2 to 6 months of age (PA1; 6-month-old). Thereafter, ALO was withdrawn from WT and MFS mice (+ inside red circles) until 9 months of age (PA1<ALO) (-inside red circles). The aortic root diameter was measured by ultrasonography at 6- and 9-month-old (brown and red boxes, respectively). N= 6-13. See also Table S6. (**B**) WT and MFS mice were treated with allopurinol following the preventive treatment (PE, 3-months-old). Thereafter, allopurinol was withdrawn from WT and MFS mice (+ inside red circles) until 6 months of age (PE<ALO) (-inside red circles). The aortic root diameter was measured by ultrasonography at 3- and 6-month-old (blue and brown boxes, respectively). N= 6-10. See also Table S7. Statistical analysis in A and B: Two-way ANOVA and Tukey post-test. ***^, +++^*P*≤0.001; *effect of genotype; ^+^effect of treatment.

**Figure S5.**
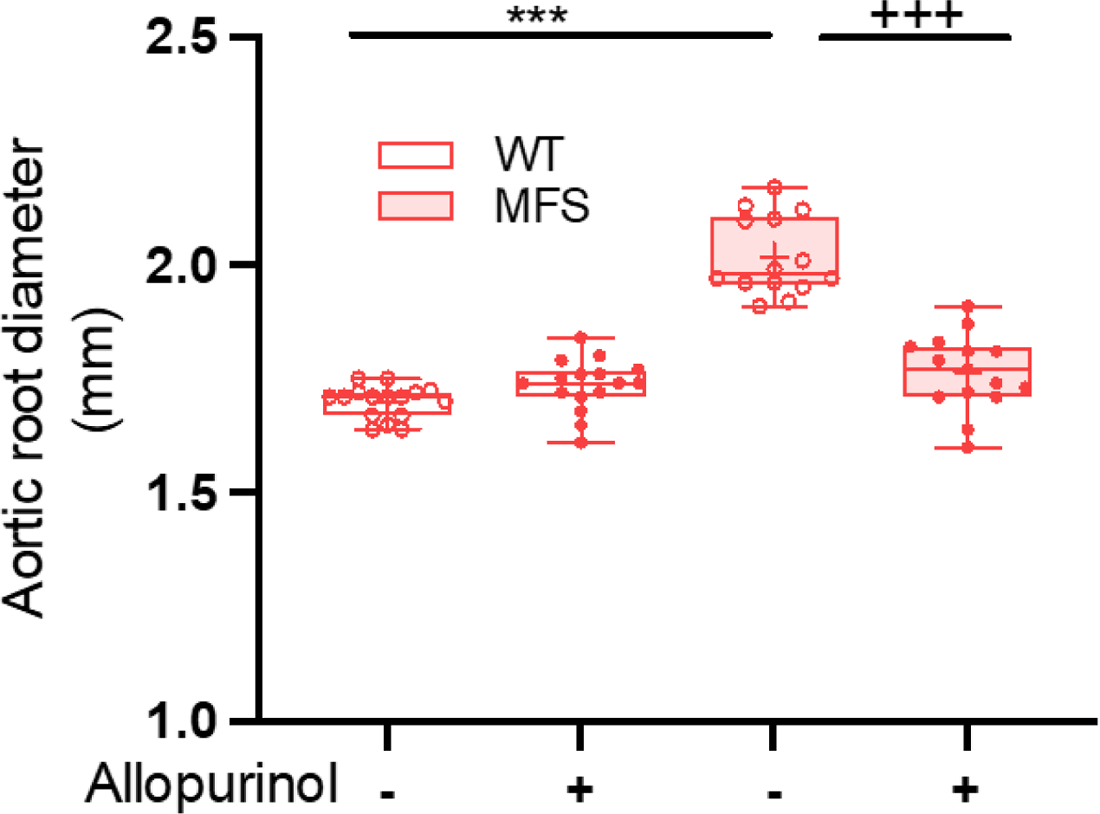
Aortic root diameter in WT and MFS mice palliatively treated with allopurinol (PA2). Data represented as boxplots. Statistical analyses: See also Table S6. Two-way ANOVA and Tukey’s post-test. ***^/+++^*P*≤0.001; *effect of genotype; ^+^effect of treatment. n=14-16.

**Figure S6.**
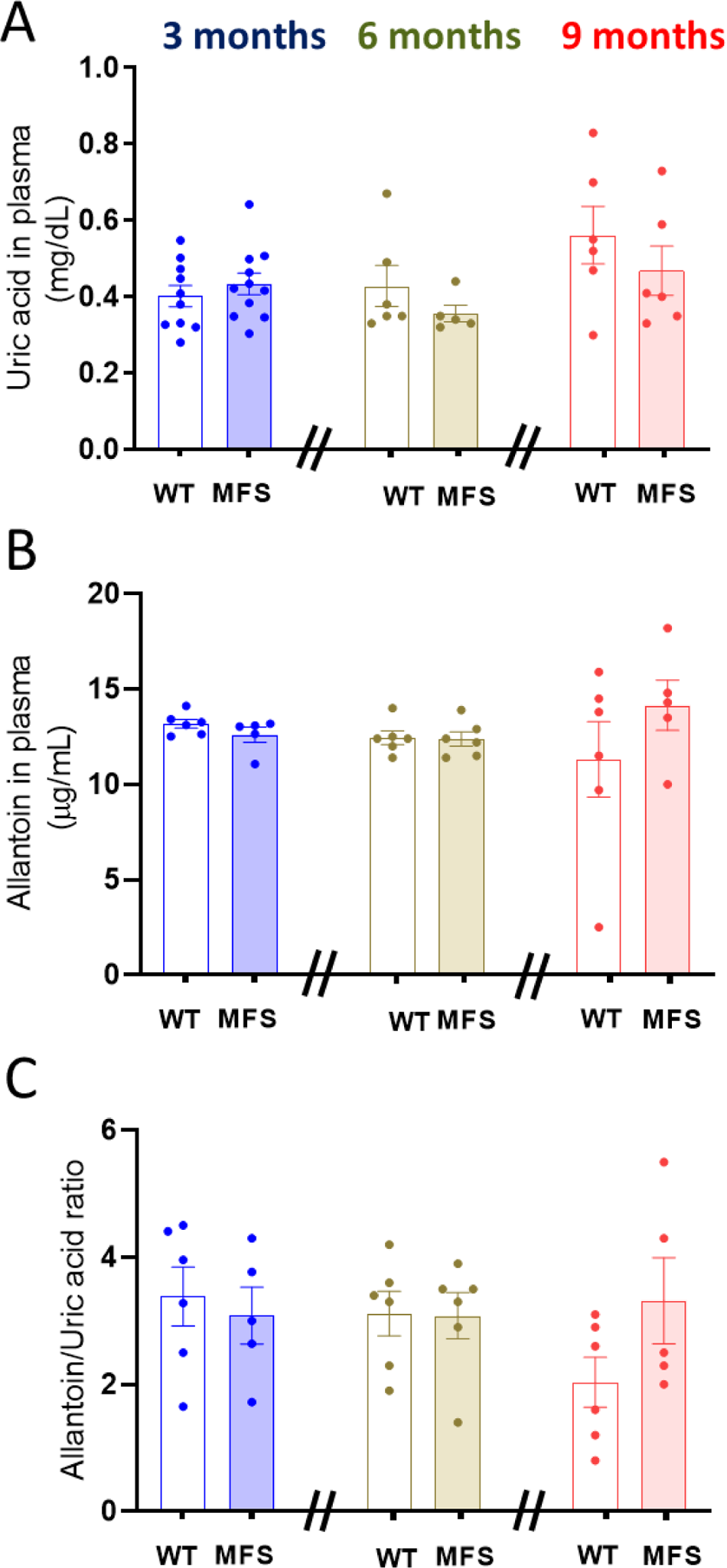
Uric acid and allantoin blood plasma levels are not altered in MFS mice. Blood plasma levels of uric acid (A), its catabolite allantoin (B) and their ratio (Allantoin/uric acid) (C) measured in WT and MFS mice of different ages (3-, 6-, and 9-month-old). Data as the mean ± SEM. Statistical analysis: Kruskal-Wallis with Dunn’s multiple comparison test. n= 5-12.

**Figure S7.**
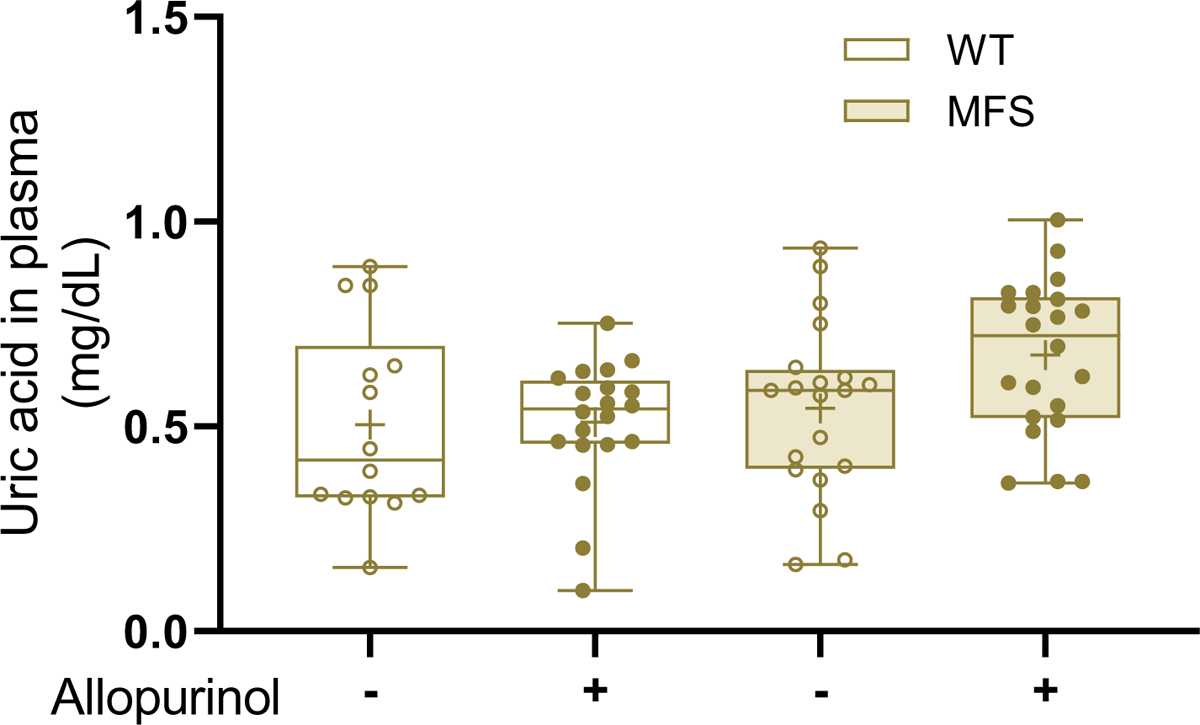
Plasma levels of uric acid do not change following allopurinol administration in WT and MFS mice. Uric acid blood plasma levels in WT and MFS mice palliatively treated with allopurinol (PA1) (n=14-22). Data represented as boxplots. Statistical test analysis: Two-way ANOVA with Tukey’s post-test.

**Figure S8.**
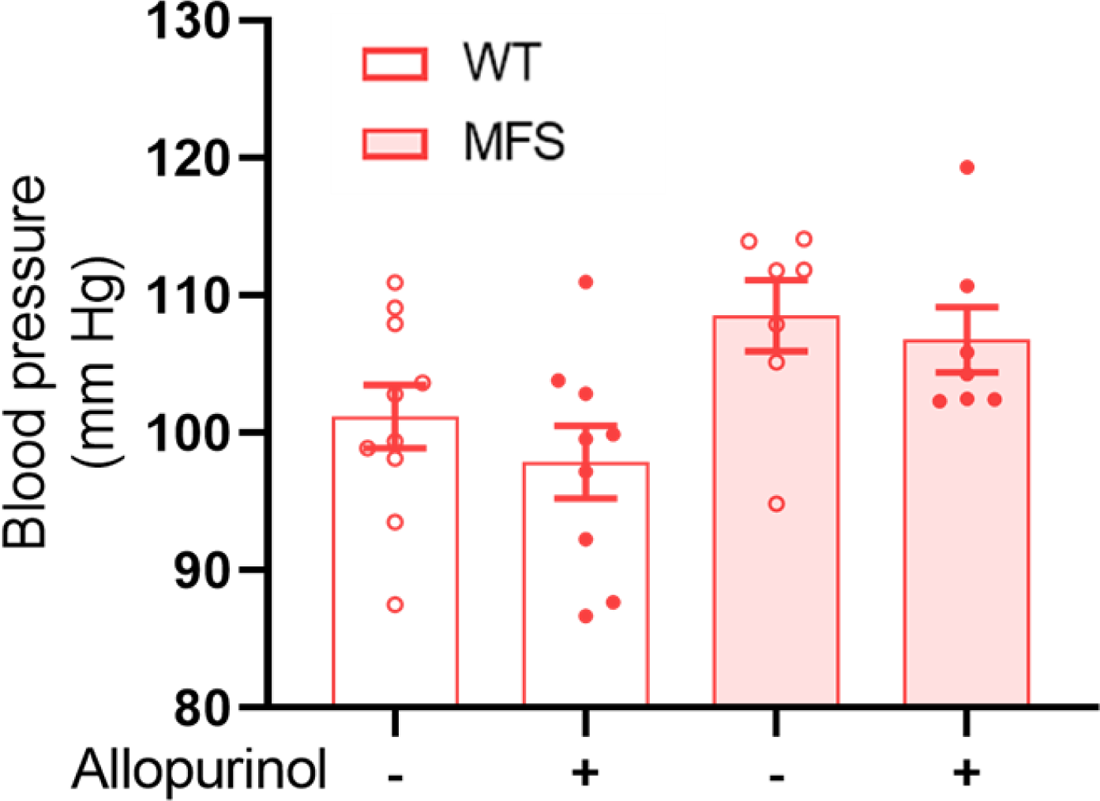
Allopurinol does not alter systolic blood pressure. Systolic blood pressure measurements in 9-month-old WT and MFS mice palliatively treated with allopurinol for 28 weeks (PA2) (n=6-10). Data as the mean ± SEM. Statistical test analysis: Kruskal-Wallis with Dunn’s multiple comparison test.

**Figure S9.**
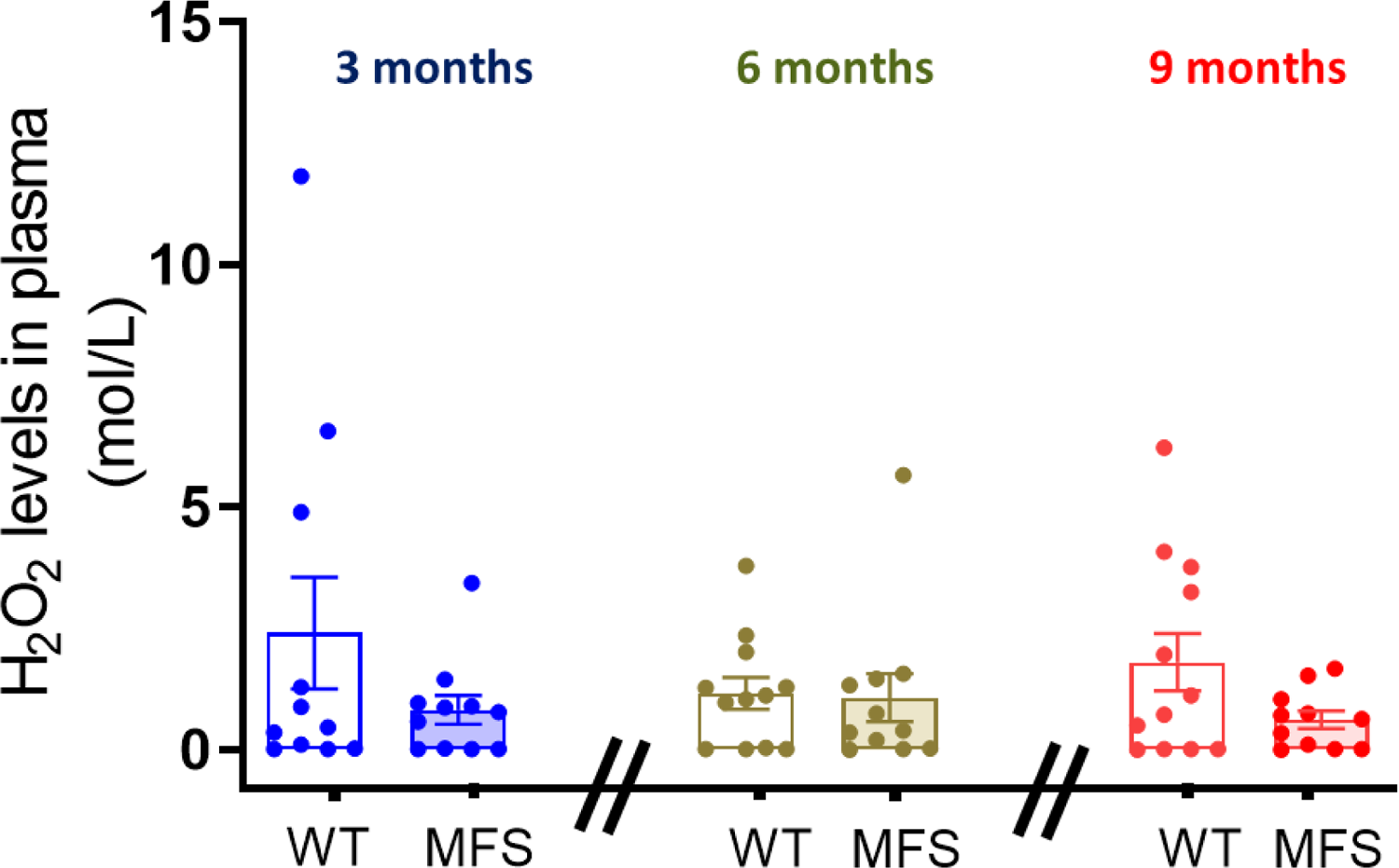
H_2_O_2_ plasma levels do not change with age in MFS mice. H_2_O_2_ levels in the blood plasma of WT and MFS of different ages (3-, 6-, and 9-monthold) (n=12-14). Data as the mean ± SEM. Statistical test analysis: Kruskal-Wallis with Dunn’s multiple comparison test.

**Figure S10.**
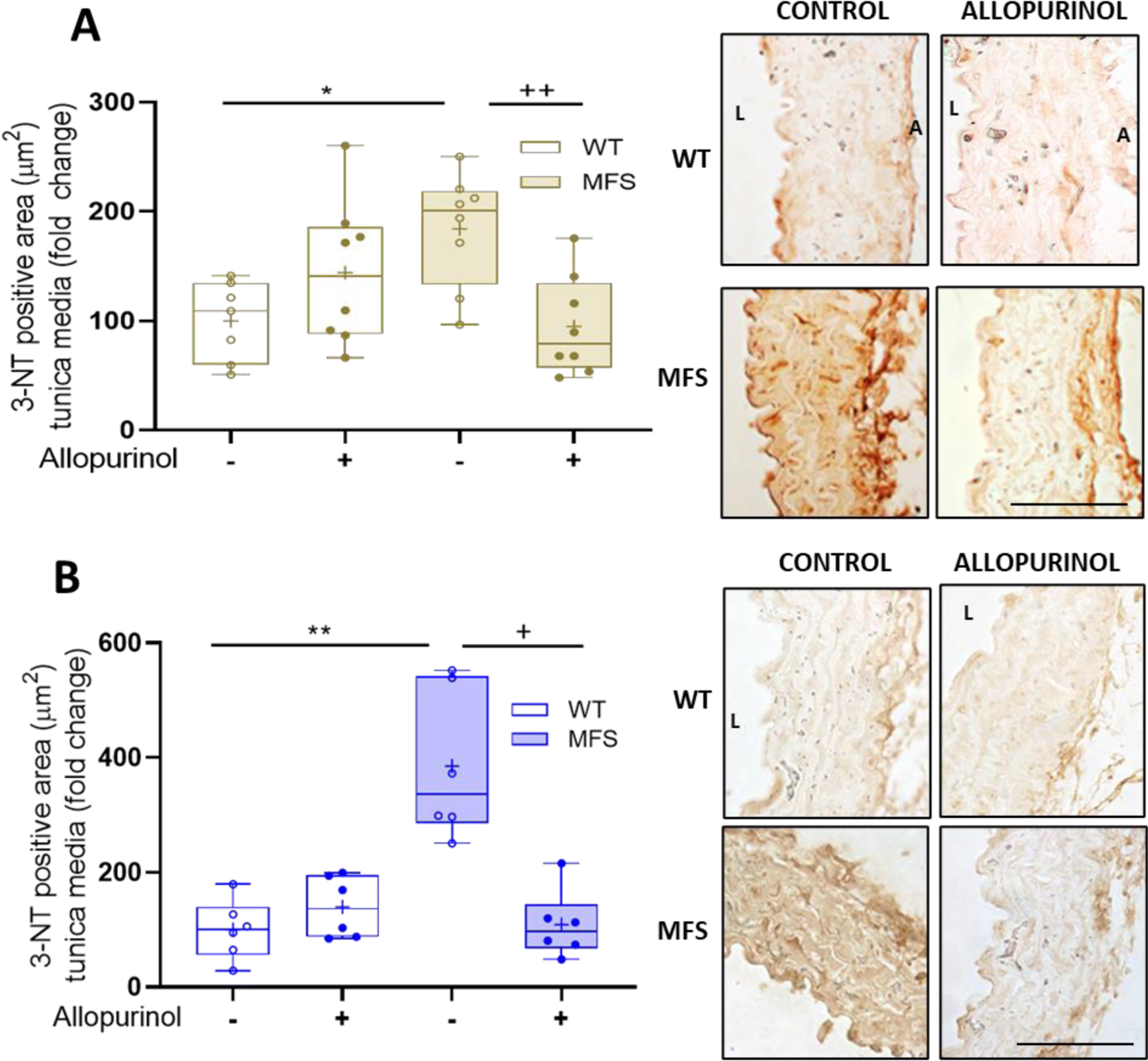
Allopurinol reduces redox stress-associated 3’-nitrotyrosine levels in the tunica media of MFS aorta. Quantitative analysis and representative images of the 3-NT levels in the aortic tunica media evidenced by immunohistochemistry with anti-3-NT antibodies after palliative (PA1) **(A)** and preventive (PE) **(B)** treatments with allopurinol in WT and MFS mice. Bar, 100 µm. Bar, 100 µm. Data represented as boxplots. Statistical test analysis: Two-way ANOVA and Tukey’s post-test (A); Kruskal-Wallis and Dunn’s multiple comparison tests (B). **^/++^*P*≤0.01 and ^*/+^*P*≤0.05; *effect of genotype; ^+^effect of treatment.

**Figure S11.**
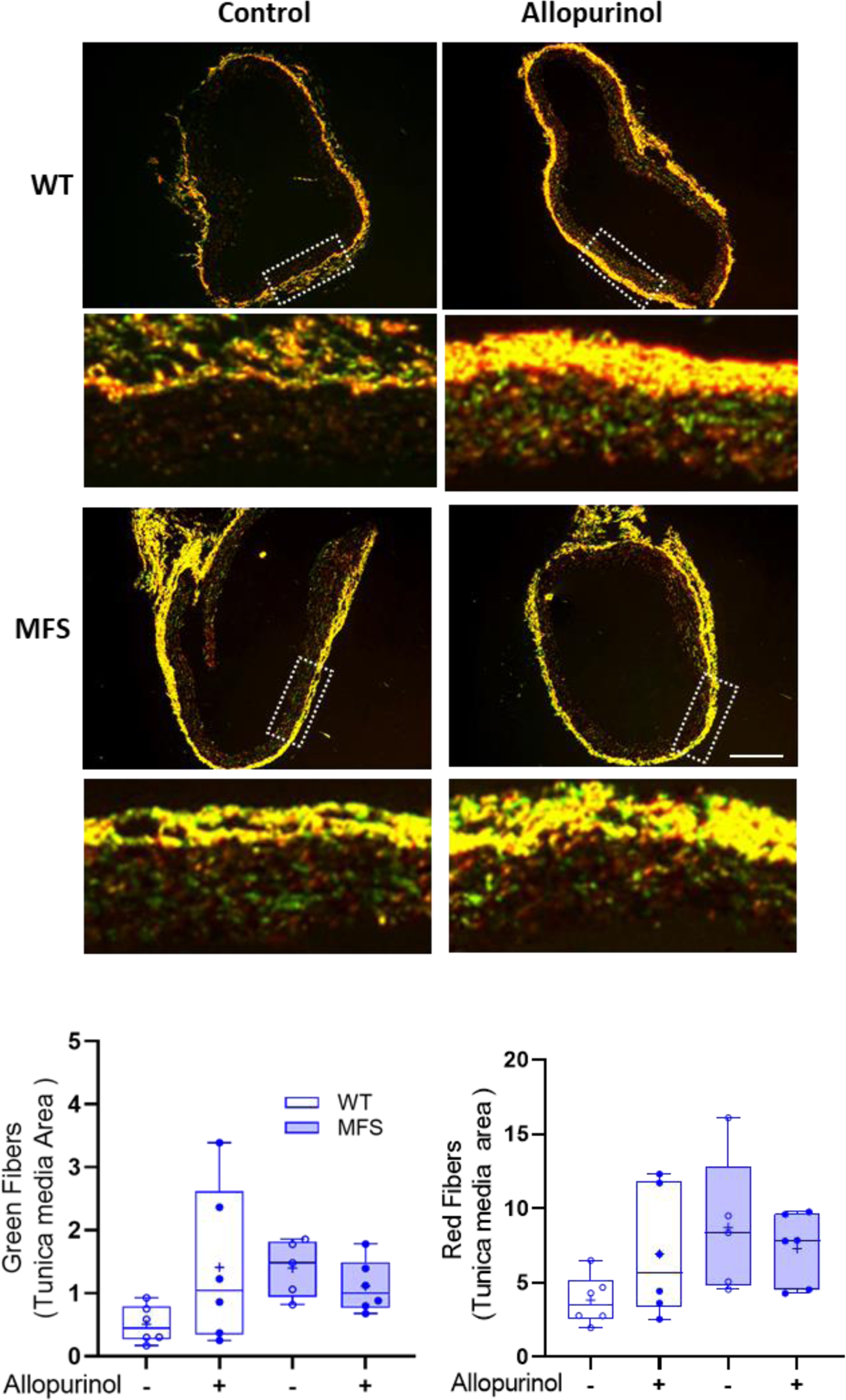
Collagen content and its maturation state in the aortic tunica media of MFS mice following the preventive treatment with allopurinol. Immature (green) and mature (red) collagen fibers of the tunicae media and adventitia of WT and MFS aortae stained with Picrosirius red visualized under the polarized microscope. WT and MFS mice were treated allopurinol in a preventive manner (PE) (n=5-6).). Representative fluorescence images of the whole aorta and enhanced indicated regions (white dashed lines). In enhanced images, the adventitia in located above and the media just below. The respective quantitative analysis of both types of collagen fibers is shown below images. Bar, 100 µm. Data represented as boxplots. Statistical test analysis: Kruskal-Wallis and Dunn’s multiple comparison tests.

### Supplemental Tables

**Table S1.**
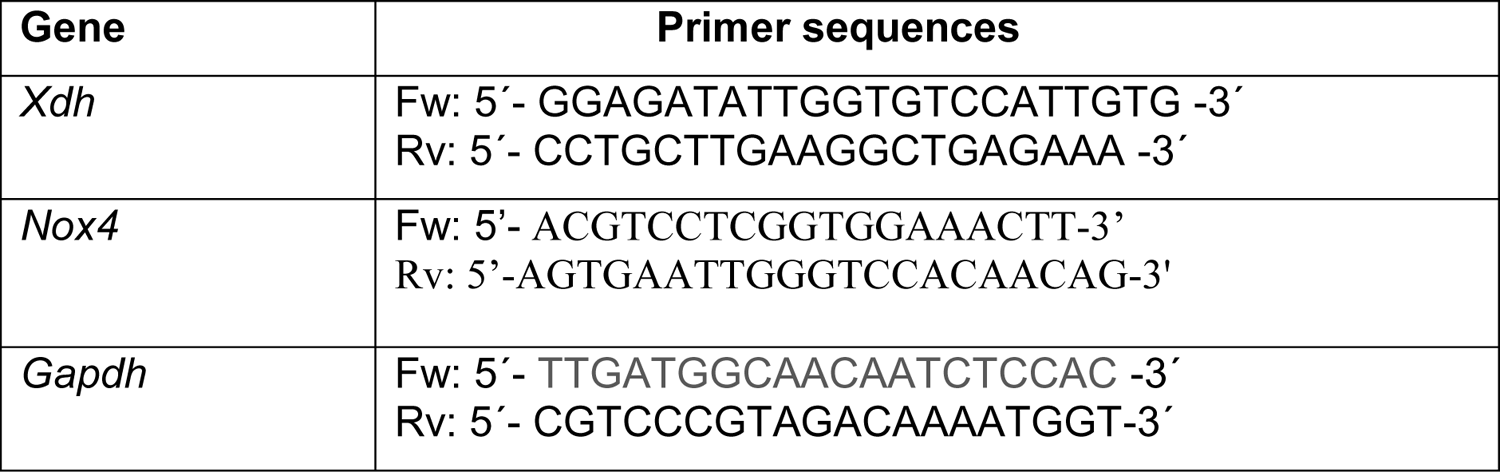
Primers used in RT-PCR analysis in MFS mice.

**Table S2.**
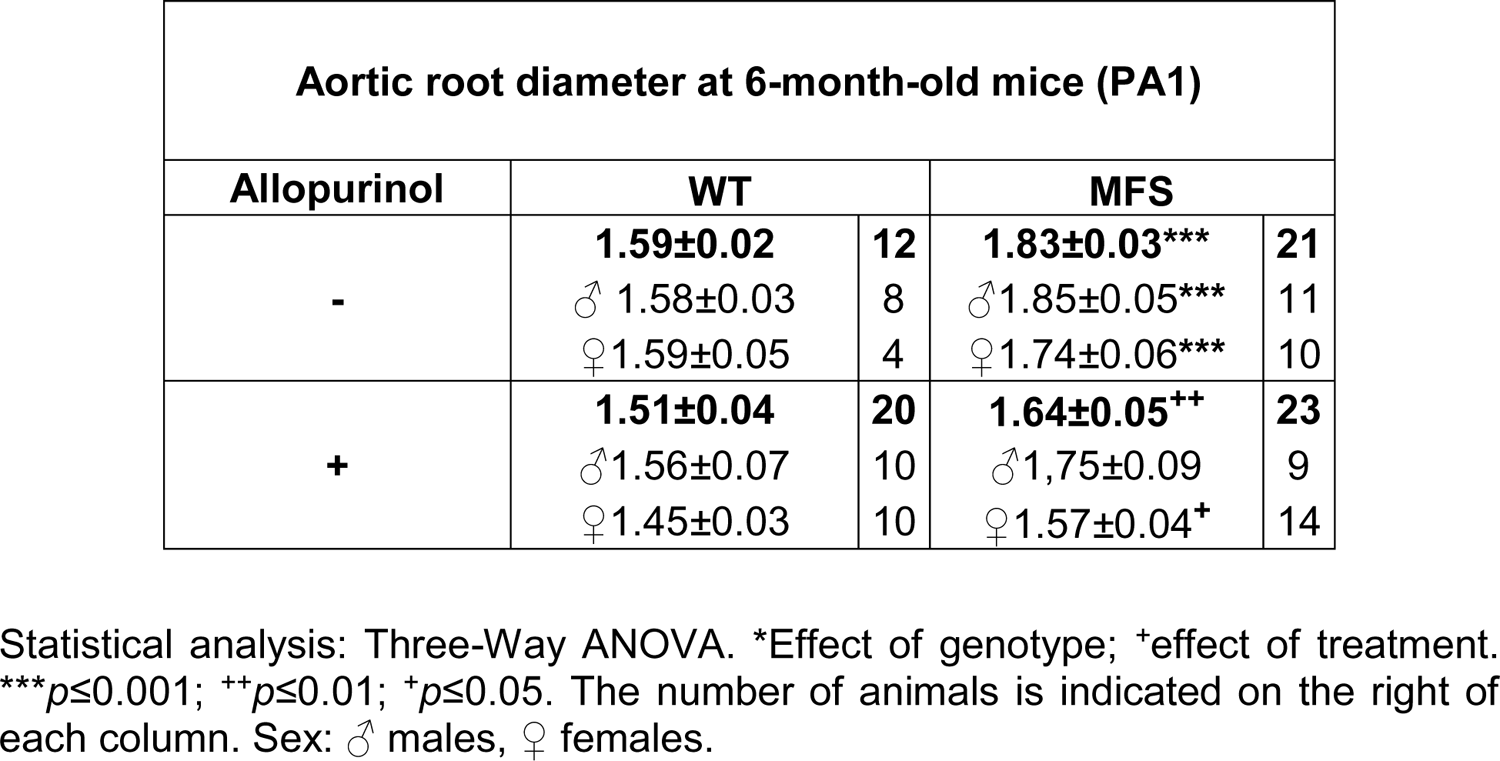
Echocardiographic values of the aortic root diameter (in mm) of WT and MFS mice subjected to palliative treatment (PA1) in the presence (+) or absence (-) of allopurinol after treatment for 4 months (from 2-to-6 month-old mice). Graphic shown in Fig. 3A. Data pooled by sex is included.

**Table S3.**
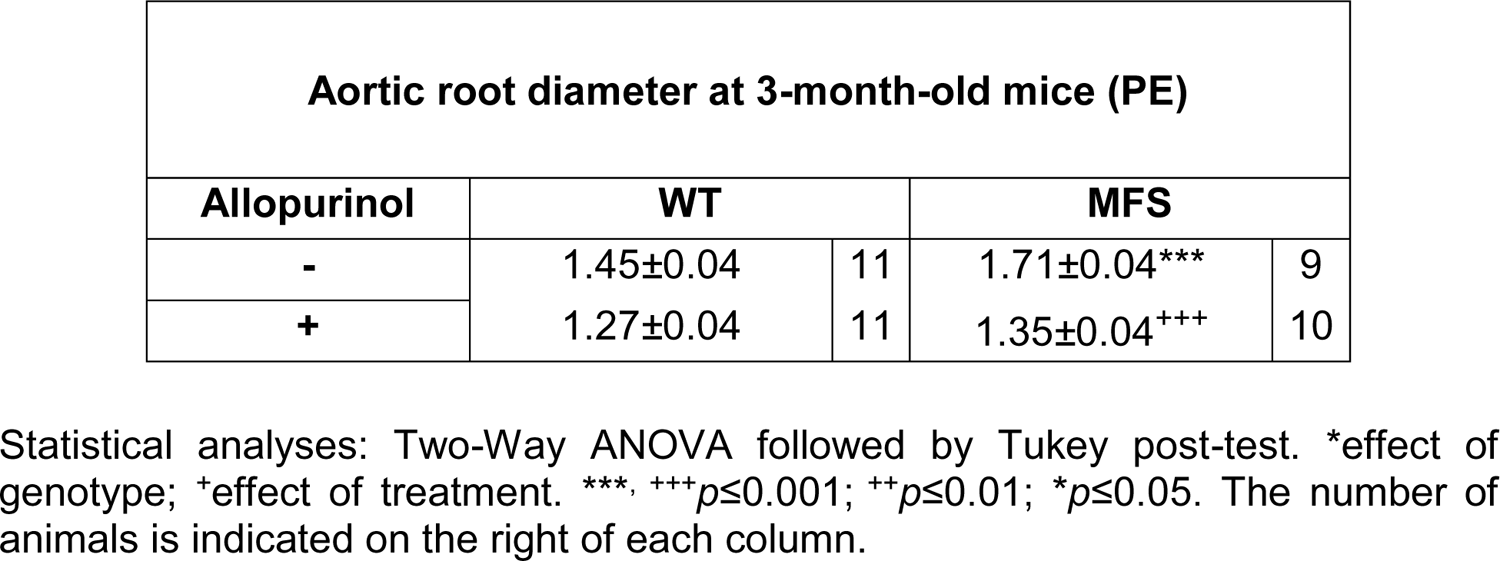
Echocardiographic values of the aortic root diameter (in mm) of WT and MFS mice subjected to preventive treatment (PE) in the presence (+) and absence (-) of allopurinol. Graphics shown in Fig. 3C.

**Table S4.** Echocardiographic values of the ascending aorta diameter (in mm) of WT and MFS mice subjected to palliative treatment (PA1) in the presence (+) or absence (-) of allopurinol after treatment for 4 months (2-to-6-month-old mice). Graphic shown in Fig. S3A.

| Ascending Aorta diameter at 6-month-old mice (PA1) |  |  |  |  |
| --- | --- | --- | --- | --- |
| Allopurinol | WT |  | MFS |  |
| - | **1.21±0.02** | **15** | **1.39±0.04\*\*\*** | **22** |
|  | ♂1.21±0.04 | 7 | ♂1.50±0.06 | 11 |
|  | ♀1.21±0.01 | 8 | ♀1.28±0.04 | 11 |
| + | **1.26±0.03** | **20** | **1.32±0.05<sup>+</sup>** | **22** |
|  | ♂1.34±0.04 | 10 | ♂1.47±0.11 | 9 |
|  | ♀1.19±0.02 | 10 | ♀1.24±0.02 | 13 |

Statistical analyses: Two-way ANOVA followed by Tukey post-test. \*Effect of genotype; <sup>+</sup>effect of treatment. \*\*\* $p \leq 0.001$ ; <sup>+</sup> $p \leq 0.05$ . The number of animals is indicated on the right of each column.

**Table S5.** Echocardiographic values of the ascending aorta diameter (in mm) of WT and MFS mice subjected to preventive treatment (PE) in the presence (+) and absence (-) of allopurinol. Graphics shown in Fig. S3B.

| Ascending aorta diameter at 3-month-old mice (PE) |  |  |  |  |
| --- | --- | --- | --- | --- |
| Allopurinol | WT |  | MFS |  |
| - | 1.10±0.16 | 11 | 1.29±0.13* | 9 |
| + | 1.17±0.16 | 11 | 1.30±0.08 | 10 |

Statistical analyses: Two-way ANOVA followed by Tukey post-test. \*Effect of genotype; <sup>+</sup>effect of treatment. \* $p \leq 0.05$ . The number of animals is indicated on the right of each column.

**Table S6.**
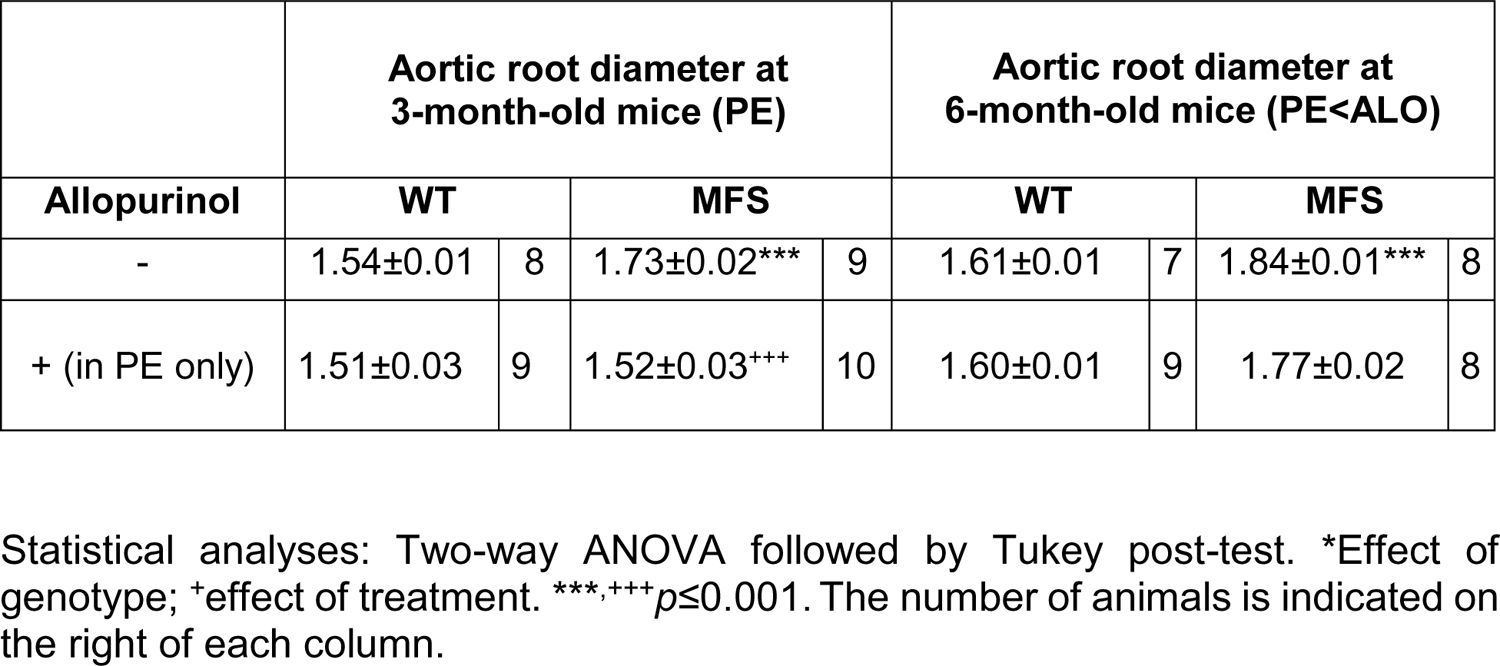
Echocardiographic values of the aortic root diameter (in mm) of WT and MFS mice subjected to preventive treatment with allopurinol from gestation until endpoint at 3-month-old /PE). Thereafter, allopurinol was withdrawn for 3 months, and mice subjected to ultrasonography (endpoint at 6-month-old/PE<ALO). Graphics shown in Fig. S4A.

**Table S7.**
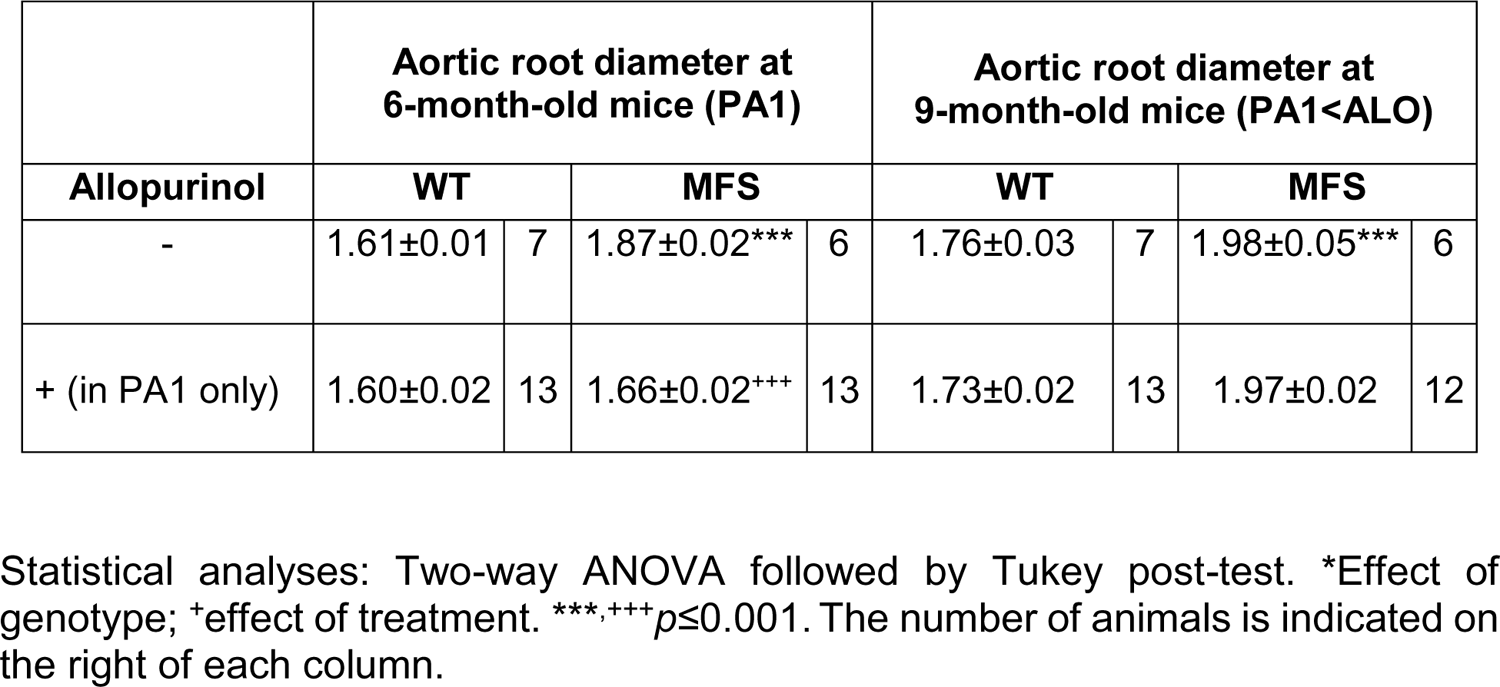
Echocardiographic values of the aortic root diameter (in mm) of WT and MFS mice subjected to palliative treatment with allopurinol (from 2-to-6-month-old/PA1). Thereafter, allopurinol was withdrawn for a period of 3 months (until 9-month-old), and mice were subjected to endpoint ultrasonography (9 month-old/PA1<ALO). Graphics shown in Fig. S4B.

**Table S8.**
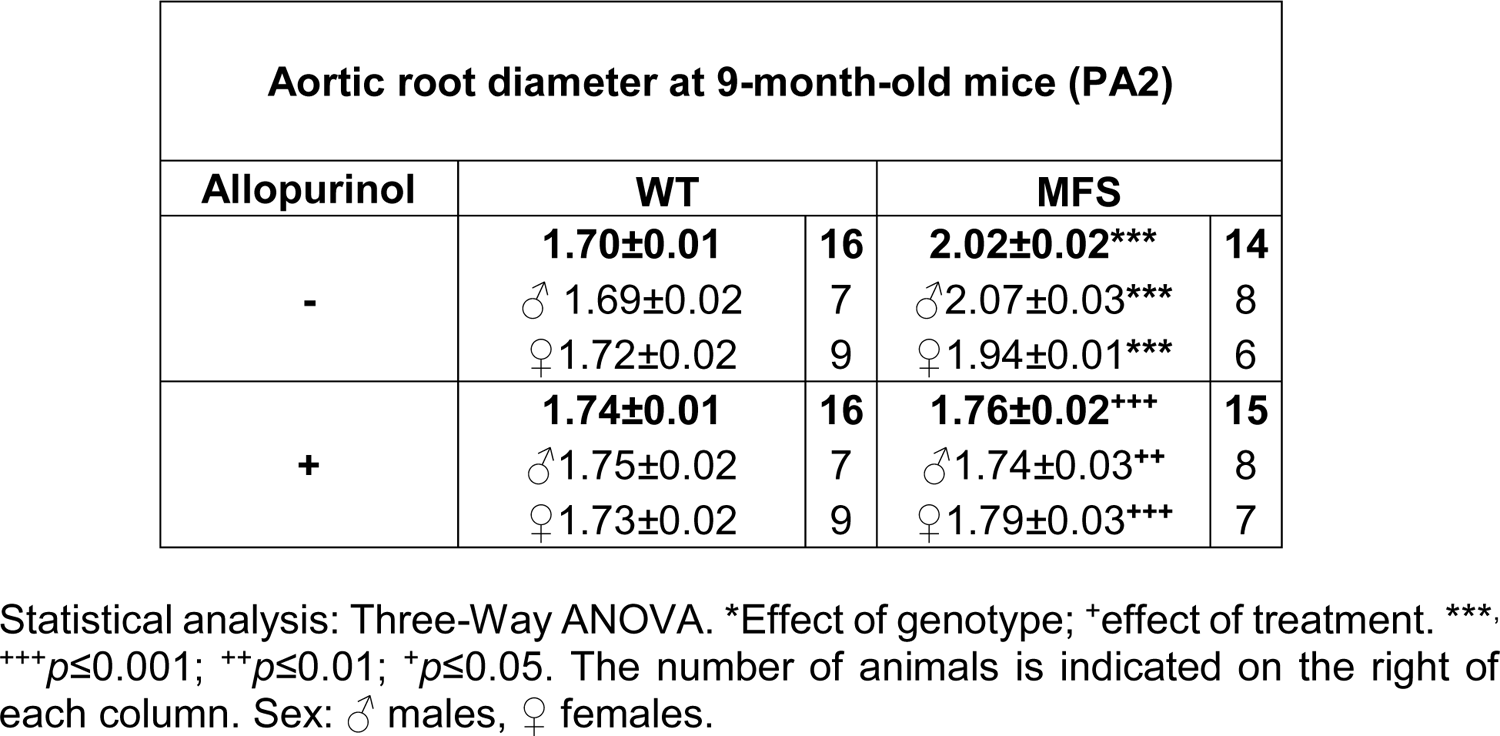
Echocardiographic values of the aortic root diameter (in mm) of WT and MFS mice subjected to palliative treatment (PA2) with allopurinol for 6 months (from 2- to 9-month-old mice). Data pooled for sex is also shown. Graphics shown in Fig. S5.

**Table S9.**
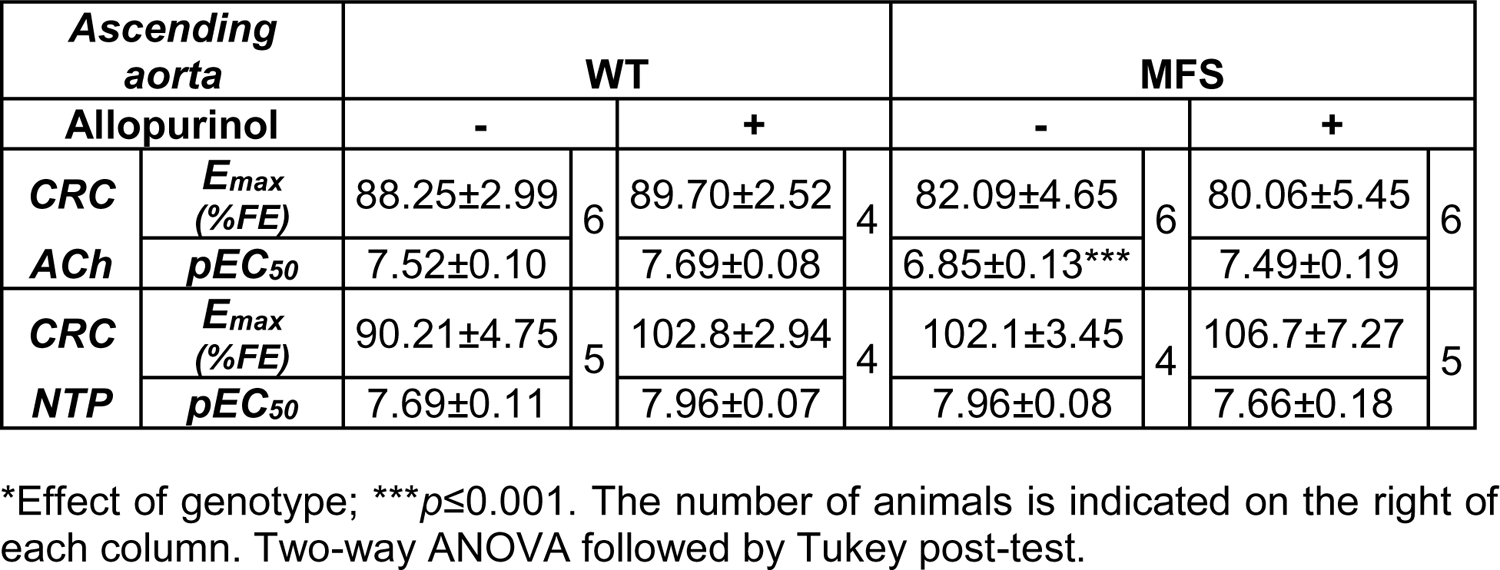
Potency (pEC_50_) and maximum response (E_max_) of the concentration– response curves (CRCs) values for ACh- and NTP-induced relaxation response (%) in the ascending aorta from WT and MFS mice (9-months-old) in the presence (+) or absence (-) of allopurinol. Graphic shown in Fig. 4D.

**Table S10.** KCl vascular tone was expressed as force units (mN). Potency (pEC_50_) and maximum response (E_max_) of the concentration-response curve (CRC) values for Phe-induced contraction response (%) in the ascending aorta of 9-month-old WT and MFS mice in the presence (+) or absence (-) of allopurinol. The number of animals is indicated on the right of each column. Graphic shown in Figs. 4B and 4E.

| **Ascending aorta** |  | **WT** |  |  |  | **MFS** |  |  |  |
| --- | --- | --- | --- | --- | --- | --- | --- | --- | --- |
| **Allopurinol** |  | **-** |  | **+** |  | **-** |  | **+** |  |
| **KCl** | $E_{max}$<br>(mN) | 4.70±0.52 | 6 | 4.56±0.80 | 5 | 4.43±0.60 | 6 | 4.32±0.49 | 6 |
| **CRC** | $E_{max}$<br>(%KCl) | 52.67±6.60 | 6 | 31.18±9.44 | 5 | 75.32±19.06 | 6 | 64.70±20.21 | 5 |
| **Phe** | $pEC_{50}$ | 6.46±0.28 | | 7.28±0.80 | | 6.25±0.61 | | 6.35±0.72 | |

**The Major Resources Table**

## SUPPLEMENTAL MATERIAL

### 1. DETAILED METHODS

G.E. has full access to the data of this study and takes responsibility for its integrity and analysis.

#### Human tissue collection, mice, and experimental designs

Healthy ascending aortic tissue was collected from heart donors via the organ donation organization at the Hospital Clínic i Provincial (Barcelona, Spain) and Hospital de Bellvitge (L’Hospitalet de Llobregat, Barcelona, Spain). The age and gender of heart donors were unknown because Spanish law protects personal information about organ donors. Ascending aortic aneurysm samples were collected from patients with Marfan syndrome (MFS) (with ages ranging from 17 to 60 years) undergoing aortic aneurysm repair surgery. All the patients in whom aortae were resected fulfilled MFS diagnostic criteria according to Ghent nosology, but no genetic information regarding putative *FBN1* mutations was available. For each patient, we obtained a 3 x 3 cm sample from two areas: the dilated zone, corresponding to the sinuses of Valsalva, and the adjacent virtually non-dilated aorta (according to the surgeon’s opinion) corresponding to the distal ascending aorta. The aortae were maintained in cold saline solution or cardioprotective solution before delivery to the laboratory.

MFS mice with a fibrillin-1 mutation *(Fbn1^C1041G/+^;* hereafter C1041G) (hereafter, MFS mice) were purchased from The Jackson Laboratory (B6.129-Fbn1tm1Hcd/J; Strain #012885/Common name: C1039G; Bar Harbor, ME 04609, USA). MFS and sex- and age-matched wild-type littermates (WT mice) were maintained in a C57BL/6J genetic background. All mice were housed according to the University of Barcelona institutional guidelines (constant room temperature at 22°C, controlled environment 12/12-hour light/dark cycle, 60% humidity and *ad libitum* access to food and water).

WT and MFS mice were administered allopurinol (hereafter ALO) (A8003, Sigma-Aldrich) diluted in drinking water (20 mg/kg/day; 125 mg/mL)^92, 93^. ALO was fully replaced each third day of treatment. We performed two experimental ALO treatment approaches: palliative (PA) and preventive (PE). For PA treatments, ALO was administered to mice of 2 months of age until 6-month-old (PA1) or 9-month-old (PA2) being a respective effective treatment of 4 and 7 months, respectively (Fig. S1). For PE treatment, ALO was administered to the pregnant WT mother, maintained after giving birth (lactation period of 25 days) and thereafter maintained in drinking water to weaned babies until three months of age. To evaluate whether ALO’s effect on aortopathy was transient or permanent, the inhibitor was withdrawn from drinking water (<ALO) for a period of three months following the PE and PA1 experimental treatments with 6- and 9-month age endpoints (PE<ALO and (PA1<ALO, respectively; Fig. S1). At each outcome time points, mice were subjected to echocardiographic analysis. Both the ascending aorta and liver were dissected and fixed for paraffin embedding for (immuno)histological studies or immersed in RNA Later (R-0901, Sigma Aldrich), frozen in liquid nitrogen and stored at −80°C for molecular tests.

#### Study approvals

Human tissues were collected with the required approval from the Institutional Clinical Review Board of Spanish clinical centers, and the patients’ written consent conformed to the ethical guidelines of the 1975 Declaration of Helsinki. Patients were informed about the use for research studies of their extracted aortic samples. All aortic tissues described in the manuscript were those obtained from Spanish Marfan patients and heart donors. Due to the Spanish Data Protection Act, we do not have access to their clinical history or personal data.

Animal care and colony maintenance conformed to the European Union (Directive 2010/63/EU) and Spanish guidelines (RD 53/2013) for the use of experimental animals. Ethical approval was obtained from the local animal ethics committee (CEEA protocol approval number 10340).

#### Echocardiography

Two-dimensional transthoracic echocardiography was performed in all animals under 1.5% inhaled isoflurane. Each animal was scanned 12–24 hours before sacrifice. Images were obtained with a 10–13 MHz phased array linear transducer (IL12i GE Healthcare, Madrid, Spain) in a Vivid Q system (GE Healthcare, Madrid, Spain). Images were recorded and later analyzed offline using commercially available software (EchoPac v.08.1.6, GE Healthcare, Madrid, Spain). Proximal aortic segments were assessed in a parasternal long-axis view. The aortic root diameter was measured from inner edge to inner edge in end-diastole at the level of the sinus of Valsalva. All echocardiographic measurements were carried out in a blinded manner by two independent investigators at two different periods, and with no knowledge of genotype or treatment.

#### Histology and histomorphometry

Paraffin-embedded tissue arrays of mice aortae from different experimental sets were cut into 5 µm sections. Elastic fiber ruptures were quantified by counting the number of large fiber breaks in tissue sections stained with Verhoeff-Van Gieson. Breaks larger than 20 µm were defined as evident large discontinuities in the normal circumferential continuity (360°) of each elastic lamina in the aortic media^45^. They were counted along the length of each elastic lamina in four different, representative images of three non-consecutive sections of the same ascending aorta. The number of sections studied for condition were usually of three, spacing 10 µm between them (2 sections). All measurements were carried out in a blinded manner by two different observers with no knowledge of genotype and treatment. Images were captured using a Leica Leitz DMRB microscope (40x oil immersion objective) equipped with a Leica DC500 camera and analyzed with Fiji Image J Analysis software.

#### Immunohistochemistry and immunofluorescence staining

For immunohistochemistry and/or immunofluorescence, paraffin-embedded aortic tissue sections (5 μm thick) were deparaffinized and rehydrated prior to unmasking the epitope. Horseradish peroxidase (HRP)-based immunohistochemistry was used to stain aortic tissue sections for XOR and 3-nitrotyrosine (3-NT). To unmask XOR epitopes, aortic tissue sections were treated with a retrieval solution (10 mM sodium citrate, 0.05% Tween, pH 6) for 30 min in the steamer at 95°C. No antigen retrieval was used for 3-NT. Next, sections were incubated for 10 min with peroxidase blocking solution (Dako Real Peroxidase-blocking solution), rinsed three times with PBS and then incubated with 1% BSA in PBS prior to overnight incubation at 4°C with the respective primary polyclonal antibodies anti-XOR (1:50; Rockland 200-4183S) or anti-3-NT (1:200; Merck Millipore 06-284). On the next day, sections were incubated with the manufacturer’s goat anti-rabbit secondary antibody solution (1:500; Abcam ab97051) for 1 h followed by the Liquid DAB+Substrate Chromogen System (Dako System HRP) for 1 min at room temperature. HRP-stained non-consecutive sections were visualized under a Leica Leitz DMRB microscope (40x immersion oil objective).

Immunofluorescence was used to stain pNRF2 in aortic sections. Sections were treated first with heat-mediated retrieval solution (1 M Tris-EDTA, 0.05% Tween, pH 9) for 30 min in the steamer at 95°C. Next, sections were incubated for 20 minutes with ammonium chloride (NH_4_Cl, 50 mM, pH 7.4) to block free aldehyde groups, followed by a permeabilization step using 0.3% Triton X-100 for 10 min and then treated with 1% BSA blocking buffer solution for 2 h prior to overnight incubation with monoclonal anti-pNRF2 (1:200; Abcam ab76026) in a humidified chamber at 4°C. On the next day, sections were rinsed with PBS, followed by 60 min incubation with the secondary antibody goat anti-rabbit Alexa 647 (1:1.000, A-21246, Invitrogen). Sections were counterstained with DAPI (1:10.000) and images were acquired using an AF6000 widefield fluorescent microscope.

For quantitative analysis of immunostainings, four areas of each ascending aorta section were quantified with Image J software. All measurements were carried out in a blinded manner by two independent investigators.

#### Uric acid, allantoin and hydrogen peroxide in blood plasma and in ascending aortic rings

Blood from mice was collected directly from the left ventricle just before the aortic tissue was dissected. Thereafter, the blood plasma was obtained by centrifugation of the blood at 3,000 rpm for 10 min at 4°C and immediately stored frozen at −80 °C. Measurement of uric acid (UA) in blood plasma was evaluated by high-performance liquid chromatography (HPLC) with ultraviolet detection. The method used for UA extraction from biological samples was an adaptation of a method previously described^94^. The plasma (100 µl) was deproteinized with 10% trichloroacetic acid. Ten µl of supernatant was injected into the HPLC system consisting of a Perkin Elmer series 200 Pump, a 717 plus Autosampler, a 2487 Dual λ absorbance detector, and a reverse-phase ODS2 column (Waters, Barcelona, Spain; 4.6 mm·200 mm, 5 µm particle size). The mobile phase was methanol/ammonium acetate 5 mM/acetonitrile (1:96:3 v/v), which was run with an isocratic regular low flow rate of 1.2 mL/min and the wavelength UV detector was set at 292 nm. UA eluted at a retention time of 2.9 minutes. Quantification was performed by external calibration. The UA detection limit in plasma was 10 ng/mL For the determination of allantoin in blood plasma, an adapted protocol was used as previously described^95^. Briefly, plasma (60 µl) was deproteinized with acetonitrile (25 µl). Samples were centrifuged (5 min, 12,000 *g*). Ten µl of supernatant was injected into the HPLC system. Separation of allantoin was performed on a Synergy Hydro-RP C-18 reversed-phase column (250 × 4.6 mm I.D., 5 m particle size) from Phenomenex (Torrance, CA, USA). Allantoin elution (at 4 min) was performed with potassium dihydrogen phosphate (10 mM, pH 2.7): acetonitrile (85:15) and ultraviolet detection (at 235 nm).

Hydrogen peroxide (H_2_O_2_) was measured both in blood plasma and in freshly dissected aortic tissue (aortic rings) utilizing a commercial assay kit (ab102500 Abcam, Cambridge, UK). Non-deproteinized blood plasma from WT and MFS mice (50 µL) and standard dilutions were mixed with 50 µL of reaction Mix (composed of 48 µL of the assay buffer plus 1 µL OxiRed Probe + 1 µL HRP for fluorometric measures) and incubated (protected from daylight) for 10 min at room temperature. In the case of the aortic tissue, two types of measurements were carried out. On the one hand, H_2_O_2_ was measured in the aorta from mice that followed the preventive treatment with ALO (*in vivo* treatment); on the other hand, H_2_O_2_ was measured in the aorta in which ALO was directly added to the assay (*in vitro* treatment) Freshly dissected ascending aorta (the adventitia was quickly removed) were cut in two portions (3-4 mm thick each) corresponding to the proximal (the half close to the heart) and distal (the half close to the aortic arch) and maintained immersed in DMEM in the culture incubator at 37°C and 5% CO_2_ until time of the assay. Subsequently, each aortic portion was incubated with the reaction mix of the kit as indicated above for the blood plasma. For the *in vitro* approach, the dissected ascending aorta was treated as above with the difference that ALO (100 µM) was added to one of the two aortic portions while the other portion received the vehicle (physiological serum/PS). Fluorometric readings were obtained at different times, reaching a plateau at 120 min. Results were normalized to the weight of the respective aortic portion. Fluorescence was measured with a Synergy fluorimeter (Ex/Em = 535/587 nm).

#### Myography tissue preparation and vascular reactivity

Segments of the ascending aorta from 9-month-old mice treated or not with ALO were dissected free of fat and placed in a cold physiological salt solution (PSS; composition in mM: NaCl 112; KCl 4.7; CaCl_2_ 2.5; KH_2_PO_4_ 1.1; MgSO_4_ 1.2; NaHCO_3_ 25 and glucose 11.1) gassed with 95% O_2_ and 5% CO_2_. Ascending aortic segments (2-3 mm) were set up on an isometric wire myograph (model 410A; Danish Myo Technology, Aarhus, Denmark) filled with PSS (37°C; 95% O_2_ and 5% CO_2_) as previously described^96^. The vessels were stretched to 6 mN as reported^93^, rinsed and allowed to equilibrate for 45 min. Tissues were then contracted twice with KCl (100 mM) every 5 min. After rinsing, vessels were left to equilibrate for an additional 30 min before starting the experiments. Vasodilatation caused by nitric oxide (NO) produced either by the endothelium itself (triggered by acetylcholine/ACh) or by the NO donor sodium nitroprusside (NTP) were determined by cumulative concentration-response curves (CRCs) of respective relaxation to ACh (10^-9^-10^-5^ M) or NTP (10^-10^-10^-5^ M) after phenylephrine (Phe; 3×10^-6^ M)-induced precontracted vessel. To study the impact of ALO on contractile responses triggered by α_1_-adrenergic stimulation, the CRCs of Phe (10^-9^ to 3×10^-5^ M)-induced contraction were evaluated. Relaxations to ACh are expressed as a percentage of Phe-precontracted level. Contractions to Phe are expressed as a percentage of the tone generated by KCl. Data from CRCs were plotted using Graph Pad Software version 8.0 (San Diego, CA, USA) with sigmoid curve fitting (variable slope) performed by non-linear regression. These curves were used to derive the values for E_max_ (the maximal relaxant response) and pEC_50_ (-log of the agonist concentration needed to produce 50% of E_max_).

#### Quantitative Real-Time PCR

Total RNA from ascending aortae was extracted using Trizol^©^ following manufacturer’s recommendations (Invitrogen, USA). RNA concentration was quantified using Nanodrop (Agilent, USA). mRNA expression levels were determined by quantitative real-time PCR (qRT-PCR) using the SYBR green detection kit. mRNA levels encoding for XOR were expressed relative to *Gadph*, which was used as the housekeeping gene. qPCR reactions were performed following the protocol guidelines of the SYBR green master mix (ThermoFisher Scientific, Waltham, MA, USA). Briefly, reactions were performed in a total volume of 10 µL, including 5 µL of SYBR green PCR master mix, 2 µM of each primer, 2 µL of nuclease-free water, and 1 µL of the previously reverse-transcribed cDNA (25 ng) template on a 368-well iCycler iQ PCR plater (Bio-Rad). All reactions were carried out in duplicate for each sample. The thermocycling profile included 45 cycles of denaturation at 95 °C for 15 seconds and annealing and elongation at 60°C for 60 seconds. Cycle threshold (Ct) values for each gene were referenced to the internal control (comparative Ct (ΔΔCt)) and converted to the linear form relative to corresponding levels in WT aortae. The primer sequences for the murine genes used in this study are shown in Table S1.

#### Fluorometric assay for measuring the enzymatic activity of xanthine dehydrogenase (XDH) and xanthine oxidase (XO) forms

XO activity was determined in WT and MFS mice from liver and aorta (in which the adventitia was previously removed) lysates using a fluorimetry-based method^97^. Part of the liver and total aorta were homogenized with five volumes per gram of tissue of 0.25 M sucrose, 10 mM DTT, 0.2 mM PMSF, 0.1 mM EDTA and 50 mM K-phosphate, pH 7.4. Homogenates were centrifuged for 30 min at 15,000 *g* and the supernatants were obtained for XO activity. XO activity was measured by calculating the slope of the increase in fluorescence after adding pterin (0.010 mmol/L), which measures the conversion of pterin to isoxanthopterin. Total activity (XO+XDH) was likewise determined after adding methylene blue (0.010 mmol/L), which replaces NAD+ as an electron acceptor. The reaction was stopped by adding allopurinol (50 µmol/L). To calibrate the fluorescence signal, the activity of a standard concentration of isoxanthopterin was measured. The extent of dehydrogenase (XDH)-to-oxidase (XO) conversion was calculated from the proportion of XO activity divided by the total activities of XDH+XO. Values were expressed as nmol/min per g of protein. The protein concentration of homogenates was determined with the Bradford assay.

#### Blood pressure measurements

Systolic blood pressure measurements were acquired in 9-month-old animals by the tail-cuff method and using the Niprem 645 non-invasive blood pressure system (Cibertec, Madrid, Spain). Mice were positioned on a heating pad and all measurements were carried out in the dark to minimize stress. All animals were habituated to the tail-cuff by daily training one week prior to the final measurements.

Then, the systolic blood pressure was recorded over the course of three days. For quantitative analysis, the mean value of three measurements per day was used for each animal. All measurements were carried out in a blinded manner with no knowledge of genotype or experimental group.

#### Collagen content measurements

Collagen content was evaluated with the PicroSirius Red Staining method. Briefly, paraffin-embedded tissue arrays of mice aortae from different experimental sets were cut into 5 µm sections. Deparaffination was performed with xylene, and rehydration in 100% ethanol three times for 5 minutes, three times in 96% ethanol for 5 min and dH_2_O for 5 min. After rehydration, samples were immersed in phophomolybdic acid 0,2% for 2 min and rinsed with distilled H_2_O. Then, samples were immersed in picrosirius red (previously prepared with picric acid (Fluka 74069) and direct red 80 (Aldrich 365548) for 2 h. Afterward, samples were rinsed with distilled H_2_O for 5 min and immersed in HCl 0,01 N for 2 min. To avoid background, unstaining was performed with ethanol 75° for 45 sec. Finally, slides were dehydrated with absolute ethanol for 5 min and fixated with xylene twice for 10 min each. Mount with DPX with coverslips. Images were captured using a Leica Leitz DMRB microscope (10x and 40x oil immersion objective) equipped with a Leica DC500 camera and analyzed with Fiji Image J Analysis software with a predesigned macro program.

#### Statistics

Data were presented either as bars showing mean ± standard error of the mean (SEM) or median±interquartile range (IQR) boxplots, in which the error bars represent minimum and maximum values, the horizontal bars and the crosses indicate median and mean values, respectively, and the extremities of the boxes indicate interquartile ranges. Firstly, normal distribution and equality of error variance data were verified with Kolmogorov-Smirnov/Shapiro Wilk tests and Levene’s test, respectively, using the IBM SPSS Statistics Base 22.0 before parametric tests were used. Differences between three or four groups were evaluated using one-way or two-way ANOVA with Tukey’s *post-hoc* test if data were normally distributed, and variances were equal or Kruskal-Wallis test with Dunn’s *post-hoc* test if data were not normally distributed. For comparison of two groups, the unpaired t-test was utilized when the data were normally distributed, and variances were equal or the Mann-Whitney U test if data did not follow a normal distribution. A value of *P*≤0.05 was considered statistically significant. Data analysis was carried out using GraphPad Prism software (version 9.1.2; GraphPad Software, La Jolla, CA). Outliers (ROUT 2%, GraphPad Prism software) were removed before analysis.

